# Endophilin-A controls recruitment, priming and fusion of neurosecretory vesicles

**DOI:** 10.1101/540864

**Authors:** Sindhuja Gowrisankaran, Vicky Steubler, Sébastien Houy, Johanna G. Peña del Castillo, Monika Gelker, Jana Kroll, Paulo S. Pinheiro, Nils Halbsgut, Nuno Raimundo, Jakob B. Sørensen, Ira Milosevic

**Affiliations:** European Neuroscience Institute (ENI), Synaptic Vesicle Dynamics Group; University Medical Center Göttingen (UMG), Göttingen, Germany; University of Copenhagen, Department for Neuroscience, Faculty of Health and Medical Science, Copenhagen, Denmark; Institute for Cellular Biochemistry, University Medical Center Göttingen (UMG), Göttingen, Germany

## Abstract

Endophilins-A are conserved endocytic adaptors with membrane curvature-sensing and - inducing properties. We show here that, independently of their role in endocytosis, endophilin-A1 and endophilin-A2 regulate exocytosis of neurosecretory vesicles. The number of neurosecretory vesicles was not altered in chromaffin cells without endophilin, yet fast capacitance and amperometry measurements revealed reduced exocytosis, smaller vesicle pools and changed fusion kinetics. Both endophilin-A1 (brain-enriched) and A2 (ubiquitous) rescued exocytic defects, but endophilin-A2 was more efficient. Distribution of neurosecretory vesicles was altered in the plasma membrane proximity, but levels and distributions of main exocytic and endocytic factors were unchanged, and slow compensatory endocytosis was not robustly affected. Endophilin’s role in exocytosis is mediated through its SH3-domain and, at least in part, interaction with intersectin, a coordinator of exocytic and endocytic traffic. Altogether, we report that endophilins-A, key endocytic proteins linked to neurodegeneration, directly regulate exocytosis by controlling vesicle recruitment, priming and fusion.

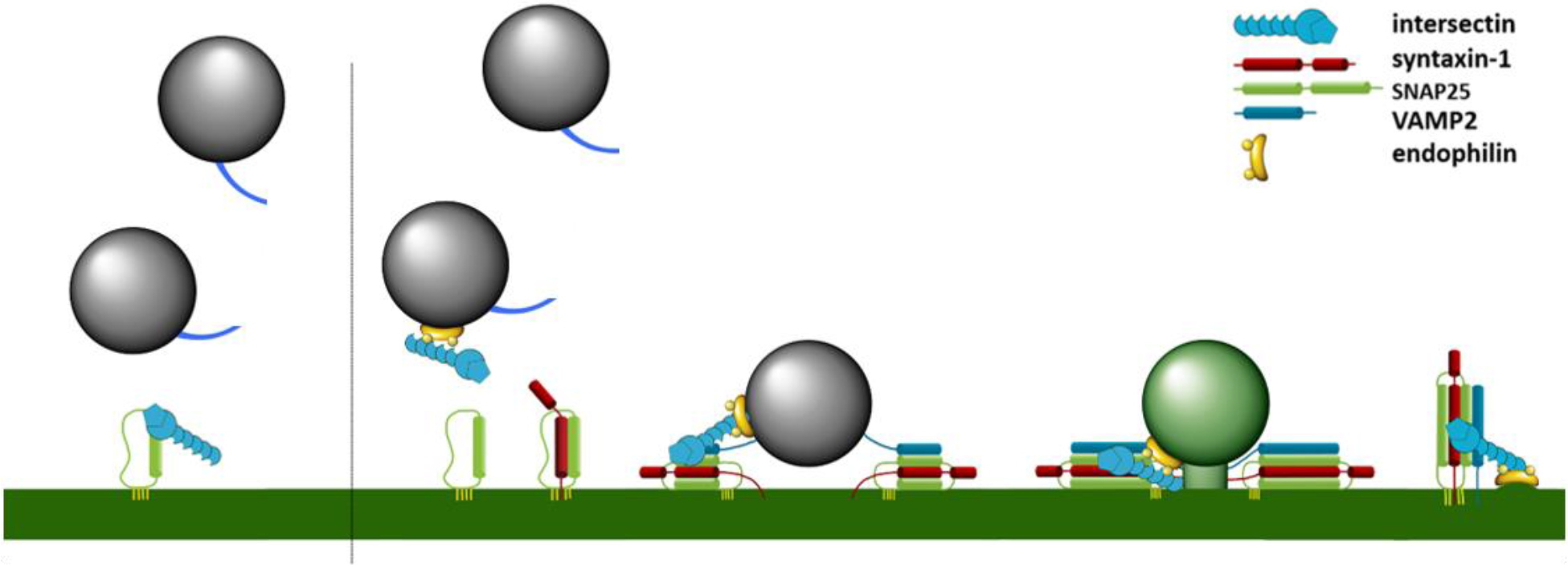

- **Recruitment, priming and fusion of secretory vesicles is controlled by endophilin**
- **Lack of endophilins alters the distribution of secretory vesicles near the PM**
- **Endophilin’s role in exocytosis is mediated through its SH3-domain**
- **Endophilin regulates intersectin localization by keeping it away from the PM**

## INTRODUCTION

Exocytosis-mediated release of vesicular content governs numerous biological events, including neurotransmission and neuromodulation, which are mandatory for brain function and survival. Following exocytosis, endocytosis rapidly retrieves the exocytosed vesicle membrane and proteins. The two processes are tightly coordinated, yet molecular mechanisms underlying such coupling are not well understood (Wu et al., 2014).

Endophilin-A (henceforth endophilin) is one of the best-characterized endocytic adaptors known to orchestrate various steps in clathrin-dependent and clathrin-independent endocytosis in mice, flies and nematodes (Ringstad et al, 1997; de Heuvel et al. 1997, Takei et al. 1999; Verstreken et al, 2002; Schuske et al, 2003; Milosevic et al., 2011; Boucrot et al., 2015; Renard et al, 2015). It contains an N-terminal Bin-Amphiphysin-Rvs (BAR)-domain that senses and induces membrane curvature, and a C-terminal SH3-domain that mediates protein interactions (de Heuvel et al., 1997; Ringstad et al., 1997). Through those two domains endophilin acts as hub of a protein network that coordinates membrane remodeling, bud constriction and scission with cargo packing, actin assembly and recruitment of factors needed for fission and/or uncoating (as reviewed in Kjaerulff et al., 2011; Milosevic et al., 2018).

Unlike invertebrates that have only one endophilin gene, vertebrates express three endophilins encoded by three genes: SH3GL2 (endophilin 1, brain-enriched), SH3GL1 (endophilin 2, ubiquitous) and SH3GL3 (endophilin 3, brain and testis-enriched). Genetic and targeted ablation studies both at invertebrate and vertebrate synapses demonstrated that the lack of functional endophilin results in defective endocytosis and an accumulation of clathrin-coated vesicles (CCVs), revealing a phenotype that is most similar to the loss of the phosphatase synaptojanin-1 (Verstreken et al, 2002; Schuske et al, 2003; Milosevic et al., 2011). Besides synaptojanin-1, other prominent endophilin interactors include dynamins, the GTPases that mediate membrane fission (Ringstad et al., 1999). Endophilin was also proposed to have a central role in clathrin-independent endocytosis (Boucrot et al, 2015; Renard et al, 2015; Simunovic et al., 2017), and, most recently, in ultrafast endocytosis (Watanabe et al., 2018).

In addition to the well-established role in endocytosis, several indications point to endophilin’s additional roles at the neuronal synapse. Specifically, endophilin interacts with synaptic vesicle (SV)-resident vesicular glutamate transporter 1 (vGLUT1; Vinatier et al., 2006; Voglmaier et al., 2006), and it was shown to undergo an association-dissociation cycle with SVs in the worm’s nerve terminal as well as to be delivered to the periactive zone by exocytosis (Bai et al., 2010). Curiously, murine neurons missing all endophilins showed a reduced number of SVs and impaired neurotransmission (Milosevic et al., 2011). Yet, the synapse relies on local SV recycling where endocytosis is coordinated with exocytosis, so it is not clear whether the impaired neurotransmission results from defective endocytosis and SV recycling, or endophilin’s role in exocytosis, or both.

Here, we show that endophilin, independently of its endocytic roles, is directly involved in the regulation of exocytosis. While exocytic and endocytic processes are tightly coupled at the synapse, in neurosecretory cells such coupling is less prominent given that the large dense-core vesicles (LDCVs) originate from the trans-Golgi network and undergo an hour(s) long maturation process before they fuse with the plasma membrane (Bader et al., 2002). Thus, we employed adrenal chromaffin cells (modified sympathetic neurons), a well-established model to study calcium-regulated exocytosis given that these cells use similar molecular machinery and release catecholamine at rates comparable to neurons in the central nervous system (Neher, 2006). After LDCVs fuse with the plasma membrane, an orchestrated process of compensatory endocytosis efficiently removes added membrane and proteins, and delivers them to a near-Golgi area, a process that takes tens of minutes (Houy et al., 2013). We found out that, like neurons, adrenal chromaffin cells contain all endophilins, thus, we employed murine chromaffin cells without endophilins (obtained from mice described in Milosevic et al., 2011) to decipher the role of endophilin in exocytosis.

## RESULTS

### Endophilin-A is enriched at the neurosecretory vesicles in chromaffin cells

Chromaffin cells contain numerous LDCVs that release their content into the blood by fast exocytosis (Neher, 2006). To check if endophilin is expressed in adrenal chromaffin cells, we looked for the presence of three endophilins’ mRNAs and proteins. Firstly, RNA was isolated from cells extracted from the center of adrenal gland (medulla) obtained from wild-type (WT) and endophilin 1, 2, 3 triple knock-out (TKO) P0 mice. All three endophilin mRNAs were detected by real-time quantitative PCR (qPCR) in the WT (endophilin 2 signal was the most prominent), but not in the TKO samples (**Figure 1A**). Next, Western blots revealed the presence of endophilin 1 and endophilin 2 in the adrenal gland homogenate from WT P0 mice, whereas these proteins were absent in the glands obtained from TKO mice (**Figure 1B**). Endophilin 3, known to be the least abundant member of the endophilin family in the brain (Milosevic et al., 2011), could not be detected in 50 μg of adrenal gland homogenate by Western blotting (**Figure 1B**), suggesting that the very small amounts of endophilin 3 mRNA detected in chromaffin cells are not efficiently translated in this system.

**Figure 1.**
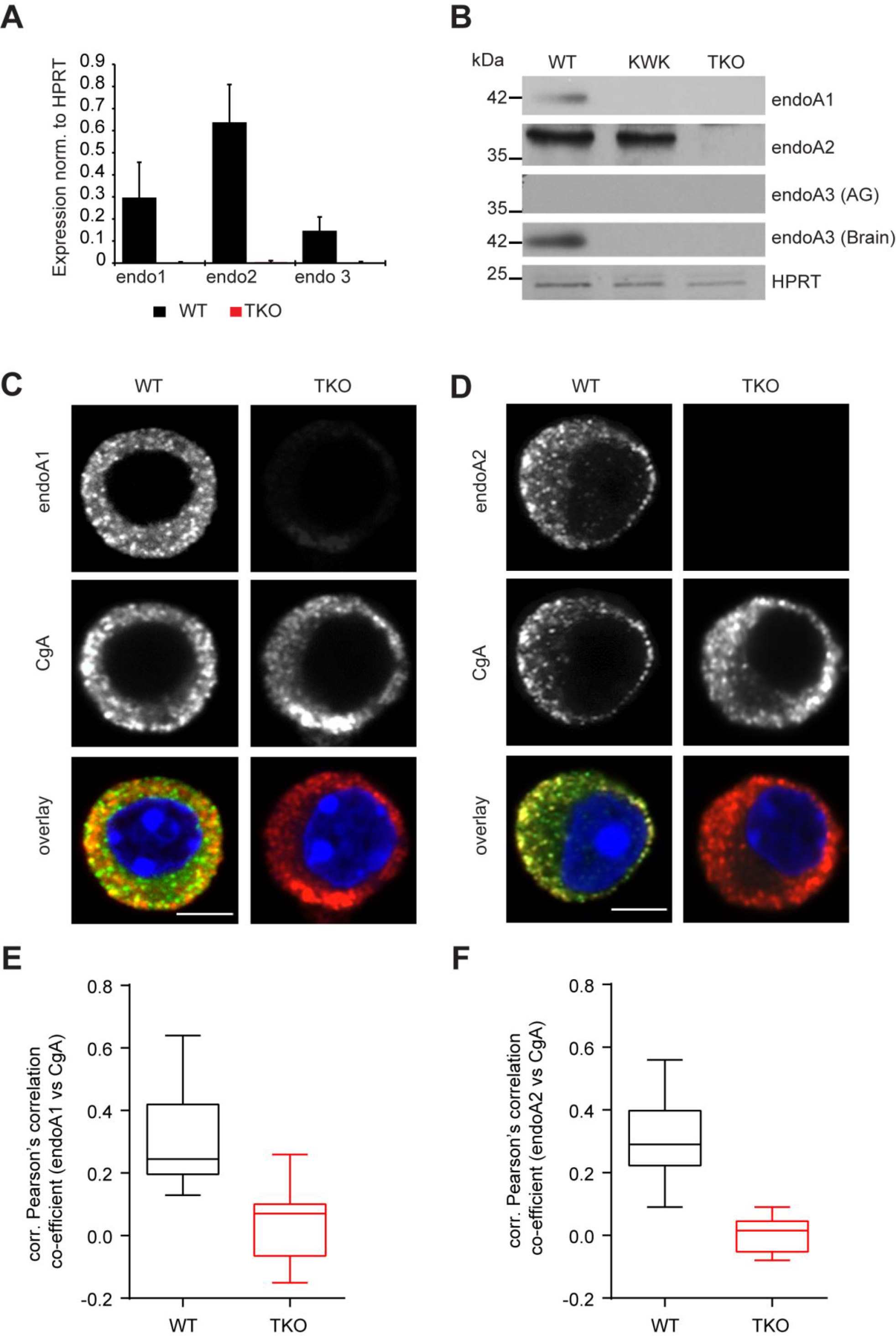
Endophilins are present in adrenal chromaffin cells and enriched at the neurosecretory vesicles. (A) Real-time PCR for endophilin 1, 2 and 3 performed on RNA isolated from the cells at the center of adrenal gland (medulla) showed the presence of all three endophilin’s mRNAs. (B) Western blot analysis of adrenal gland lysate blotted with anti-endophilin 1, 2 and 3 antibodies. Endophilin 3 could not be detected in the adrenal gland homogenate, although it could be detected in the same amount of WT brain sample. (C-D) Confocal images of WT and endophilin TKO mouse chromaffin cell stained with endophilin 1 (C) and endophilin 2 (D), and co-labeled with chromogranin-A (CgA), a LDCV marker. The images show an optical section through the cell’s equatorial plane. The endophilin antibodies used were characterized as specific since little to none signal could be detected in endophilin TKO cells. (E-F) Colocalization analysis of CgA with endophilin 1 and endophilin 2, respectively, was performed in WT and endophilin TKO cells as detailed in Methods (note that the accidental colocalization was subtracted, resulting in negative correlation in some endophilin TKO cells).

The studies of native endophilins’ distribution in mammalian cells are hindered by the lack of specific antibodies for immunostainings. Based on the overexpression studies, endophilins are primarily cytosolic proteins that associate with membranes in various cells (Ringstad et al., 2001; Perera et al., 2006; Milosevic et al, 2011; Murdoch et al, 2016). To check for the distribution of endophilins in chromaffin cells, we generated two new anti-endophilin antibodies and tested their specificity on knock-out cells and tissues. These antibodies, termed endo-A1gp and endo-A2rab respectively, gave almost no staining in endophilin TKO cells (**Figure 1C** and **1D,** right panels) and revealed that the majority of endophilin 1 and 2 was cytosolic (**Figure 1C** and **1D,** left panels). Yet, a part of endophilin 1 and endophilin 2-specific signal was punctate and reminiscent of signal obtained with the LDCV makers, e.g. chromogranin-A (CgA) (**Figure 1C** and **1D**). When chromaffin cells were co-immunostained for CgA and endophilin 1 (or endophilin 2, respectively), a significant colocalization between CgA and endophilins was detected (**Figure 1E** and **1F;** note that the colocalization values were corrected for an accidental colocalization given the high abundance of both signals).

To test if endophilin can indeed associate with the secretory vesicles, we purified LDCVs from the medulla of bovine adrenal glands by adapting the published protocol (**Suppl Fig S1A**; Park et al., 2012; mouse adrenals glands are small, 1-2 mm in size, and they did not provide sufficient amount of starting material), and we isolated SVs from the mouse brains (Farsi et al., 2018). Immunoblotting revealed that endophilin 1 was present in the purified LDCV as well as in the purified SV samples (**Suppl Fig S1B;** note that endophilin 1 was detected in adrenal gland homogenate in **Figure 1B**, but not here since the lower sample amounts were loaded to avoid Western blot signal saturation).

Having specific antibodies against endophilins 1 and 2 for immunostaining now available, we inspected the distribution of endophilin 1 and 2 in chromaffin cells upon stimulation (by 59 mM potassium solution). Interestingly, a mild enrichment of endophilin 1 and endophilin 2-specific signals was detected near/at the plasma membrane upon stimulation (**Suppl Fig S1C-D,** graphs show the line intensity profiles through chromaffin cells).

In sum, a fraction of endophilin-specific signal in chromaffin cells colocalized with LDCVs and translocated to the plasma membrane upon stimulation. LDCVs are derived from Golgi and undergo a long maturation process before their release, thus the presence of endophilin on LDCVs is unexpected, in particular in the light of two decades long research on this protein family. The function(s) of endophilin associated with the neurosecretory vesicles and in exocytosis are entirely unknown.

### Endophilin-A promotes exocytosis in chromaffin cells

To check whether endophilins have a role in exocytosis, we performed fast electrophysiological and electrochemical measurements that combined membrane capacitance and amperometry recordings on chromaffin cells of endophilin TKO mice and corresponding littermate control (endophilin A1^−/−^-A2^+/+^-A3^−/−^, henceforth endophilin KOWTKO or KWK; note that a direct comparison to wild-type, WTWTWT, littermates was not possible since the strain was knockout for endophilin A1 and A3). The chromaffin cells were isolated from adrenal glands at P0 (for experimentation, the cells were used between 2-4 days-in-vitro, DIV). Fast capacitance and amperometry measurements were performed as follows: each cell was loaded (via a patch pipette) with the photo-labile Ca^2+^-chelator nitrophenyl-EGTA and two Ca^2+^-dyes (Fura-4F and Furaptra) to enable accurate measurements of intracellular calcium concentrations ([Ca^2+^]_i_) during the whole experiment. Photolysis of caged Ca^2+^ compound increased [Ca^2+^]_i_ from several hundred nM to above 10 μM, resulting in a robust exocytosis, which was assayed by an increase in membrane capacitance and amperometric current, one cell at a time. The measured increase in membrane capacitance is proportional to the change in chromaffin cell plasma membrane surface area, while simultaneous amperometric recordings allowed the measurements of secreted catecholamine during the exocytic events (Milosevic et al., 2005; Nagy et al., 2005; Neher 2006). Remarkably, both capacitance and amperometry measurements from endophilin TKO chromaffin cells revealed reduced exocytosis (340±36 fF) when compared to the control littermate (501±30 fF for KOWTKO) after the first stimulation (**Figure 2A-B**). We noted that exocytosis from endophilin KOWTKO cells was comparable to the exocytosis from wild-type cells originating from several different C57/BL6-based strains that have been recorded over the years in different laboratories (e.g. Sørensen et al., 2003; Borisovska et al., 2005; Liu et al., 2008; Schonn et al., 2008; Pinheiro et al., 2014; Kedar et al., 2015; Man et al., 2015). Analysis of capacitance measurements revealed that the exocytic burst (189±25 fF in TKO vs. 299±22 fF in KOWTKO, **Figure 2C**) and the sustained component (151±16 fF TKO vs. 202±14 fF in KOWTKO; **Figure 2D**) were reduced. Further analysis of the exocytic burst revealed that both readily releasable pool (RRP) and slowly releasable pool (SRP) sizes (the amplitudes of the exponential fits) were significantly smaller (78±15 fF vs. 142±13 fF and 54±10 fF vs. 91±11 fF, respectively**; Figure 2E)**. The fusion kinetics of the RRP vesicles from the TKO cells (time constants of the exponential fits) was sped up (9.0±1.0 ms vs. 12.1±1.0 ms; **Figure 2F).** Curiously, we found that the RRP time constant correlated with the RRP size (**Suppl Fig S2**; also noted in synaptotagmin-7 KO, Schonn et al., 2008), suggesting that the faster fusion kinetics and lower amplitude may not be two phenotypes, but one. The kinetic parameter of the SRP was not significantly altered, although on average it was slower in the TKOs (**Figure 2G**). Altogether, these data indicate that endophilin controls the size of releasable pools and the rate of RRP vesicle fusion. Similar results were found upon a second stimulation applied 100s after a first stimulation (**Figure 2H-K**; small exocytic responses prevented reliable exponential fitting so the vesicle pool and kinetic analyses could not be performed). Interestingly, the ratio between second and first burst was not changed (**Figure 2L**), suggesting that vesicle pools could be efficiently refilled between two stimuli.

**Figure 2.**
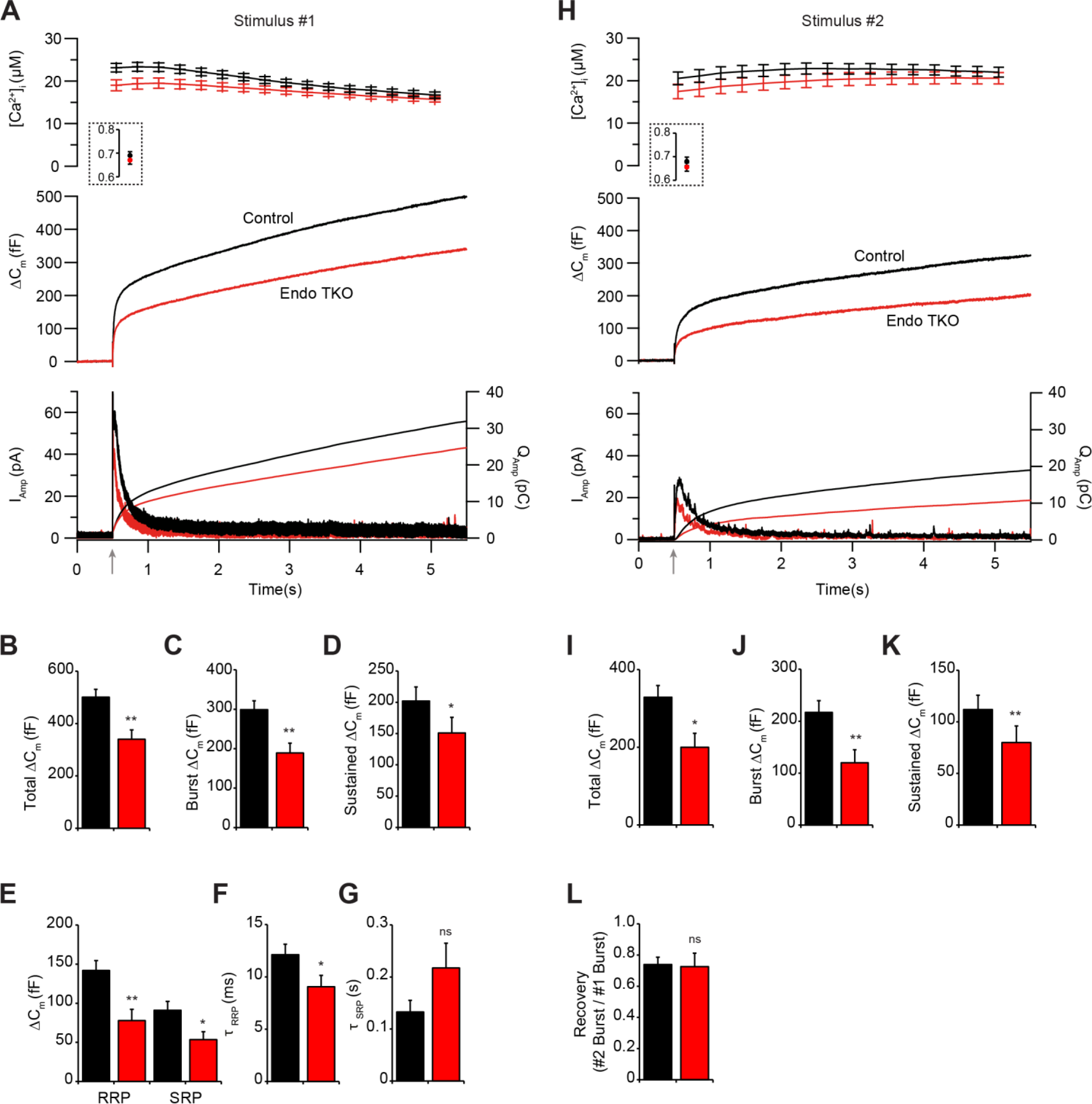
Lack of endophilins reduces exocytosis in mouse chromaffin cells. (A) Exocytosis induced by the UV-flash photolysis of caged calcium (stimulus #1, at arrow) was reduced in endophilin TKO chromaffin cells (red traces) compared to littermate KOWTKO control cells (black traces). (A) Top: intracellular calcium level increase induced by flash photolysis at 0.5s (at arrow). The inset shows the pre-flash calcium levels in the cells. Middle: averaged traces of membrane capacitance changes upon Ca^2+^-induced exocytosis. Bottom: mean amperometric current (left axis) and cumulative charge (right axis). (B-D) Analysis of capacitance traces (30 control cells - black bars, 29 TKO cells - red bars) revealed an overall reduction of exocytosed vesicles. Exponential fitting of the capacitance traces revealed changes in burst (exocytosis within the first 1s; C) and sustained phase of release (D). (E) Both RRP and SRP components of the burst phase were significantly reduced in endophilin TKO cells. (F) Fusion kinetics of RRP vesicles was faster in the TKO cells (unpaired t-test). (G) Although on average slower, fusion kinetics constant of SRP was not significantly changed in endophilin TKO cells (unpaired t-test). (H-K) Exocytosis induced by a second stimulus, elicited 100s after the first stimulus, showed similar reduction in total vesicle exocytosed as well as burst and sustained components of the release (the smaller exocytic responses prevented exponential kinetic analysis). Panels in (H) are arranged as detailed in (A). (L) Measure of recovery, calculated as the ratio of burst secretion of the second over the first stimulus was not significantly altered.

To verify that the effect of lack of endophilin on exocytosis is specific, and to test for the contribution of individual endophilins to the chromaffin cell secretion, we expressed full length endophilin 1, or endophilin 2 respectively, in endophilin TKO cells using a bicistronic lentiviral system, and performed electrophysiological measurements as before. The lentiviral expression system was verified in the HEK-293 cells (**Suppl Fig S3A).** The co-expression of enhanced green fluorescent protein (EGFP) through the IRES system was used as a marker of expression in each recorded cell, and it indicates that proteins were expressed to similar levels (**Suppl Fig S3B)**. Remarkably, both endophilin 1 and endophilin 2 were able to rescue exocytosis in TKO cells (**Figure 3A-B**; the rescue with endophilin 2 was indistinguishable from endophilin KOWTKO cells). Both burst and sustained exocytic components were efficiently rescued by either endophilins (burst: 221±28 fF for endophilin 1 and 292±43 fF for endophilin 2 vs. 137±15 fF measured in endophilin TKO; sustained: 190±23 fF for endophilin 1 and 178±20 fF for endophilin 2 vs. 92±24 fF in endophilin TKO; **Figure 3C-D**), as well as the RRP (**Figure 3E**) and the fusion kinetics of the RRP (**Figure 3F**). Although the kinetics of SRP was faster when either endophilin 1 or 2 were expressed, the difference was not significant (**Figure 3G**). Similar results were noted upon a second stimulation (**Suppl Fig S3C-F**). Notably, the releasable pools were recovered efficiently between two stimuli that were 100s apart, as revealed by an unchanged ratio between second and first burst (**Suppl Fig S3G**). Altogether, these data reveal that the effect of endophilin on exocytosis was specific, and that the expression of either endophilin 1 or 2 was sufficient to support exocytosis in chromaffin cells.

**Figure 3.**
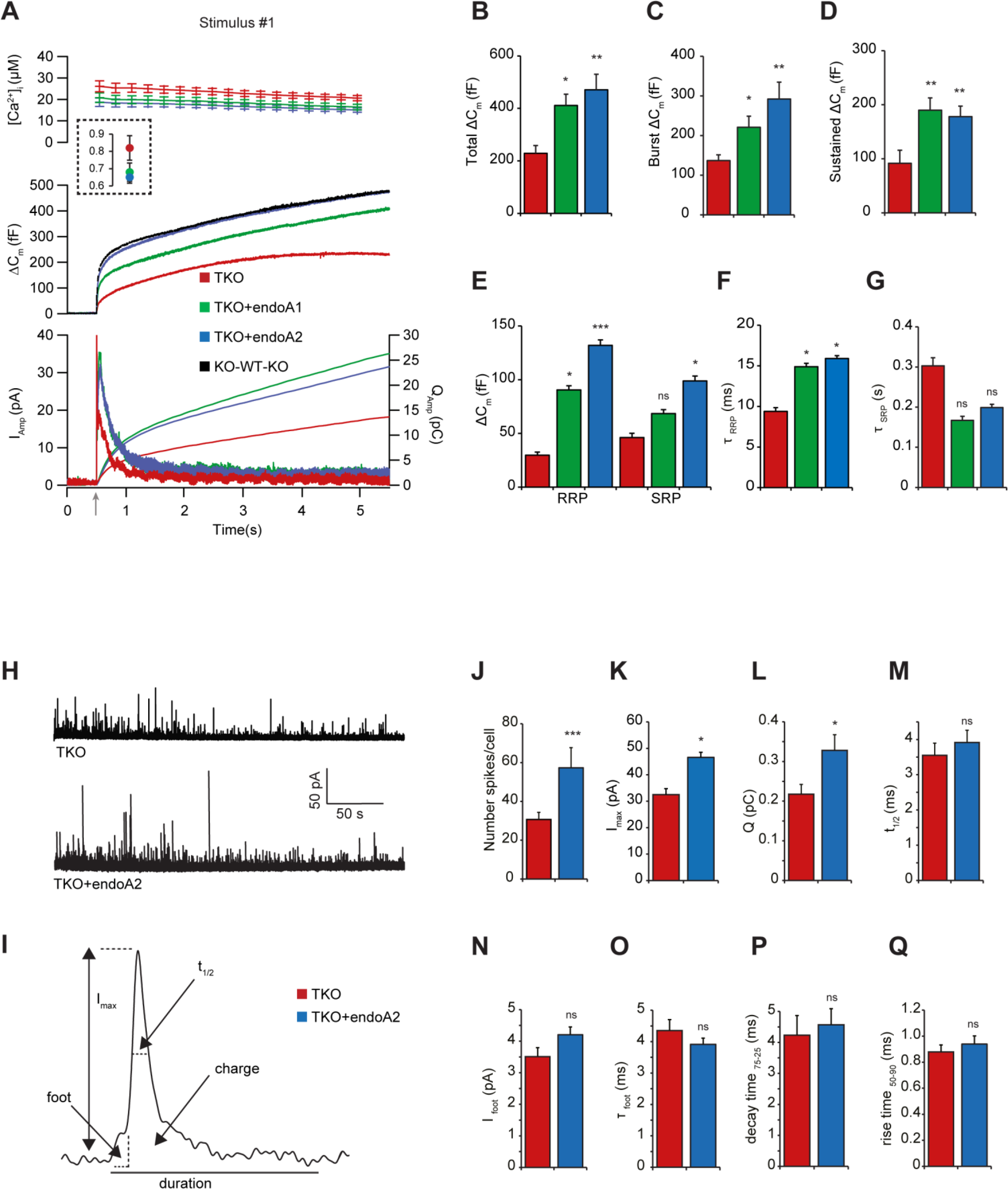
Expression of endophilin 1 and endophilin 2 rescued exocytosis in endophilin TKO cells. (A-B) Expression of endophilin 1 and endophilin 2 using bicistronic lentiviral system rescued the exocytic defects seen in endophilin TKO cells. Panel arranged as in Figure 2A, with 3 groups: endophilin TKO (red traces; mean of 24 cells), endophilin TKO + endophilin 1 (green traces; mean of 23 cells) and endophilin TKO + endophilin 2 (blue traces; mean of 25 cells). Endophilin KOWTKO data from Figure 2A (black trace; mean of 30 cells) are superimposed. Note that both endophilin 1 and endophilin 2 can rescue exocytosis, rescue with endophilin 2 is indistinguishable from control KOWTKO cells. (C-E) Burst and sustained component, as well as RRP size, were rescued upon expression of endophilin 1 and 2, respectively. (F-G) The altered kinetics of RRP in TKO was rescued (F), while time constant of SRP was not significantly changed (G) upon expression of endophilin 1 and 2, respectively (Kruskal-Wallis test with Dunn’s multiple comparison test). (H) Exemplary traces from amperometric recordings of endophilin TKO and endophilin TKO expressing endophilin 2. (I) Schematic of analyzed amperometric spike parameters. (J-Q) Amperometry analysis reveals problems in vesicle fusion: number of fusion events per cell (I), single spike amplitude (J) and charge (K) were significantly decreased in endophilin TKO cells, while the rise time (P) and foot properties (N-Q) were unaltered (Mann-Whitney test).

To inspect whether chromaffin cells without endophilin show qualitative changes in vesicle fusion, we examined the properties of single amperometric events from endophilin TKO cells and TKO cells expressing endophilin 2. Here, secretion was elicited by loading chromaffin cells in a whole-cell mode with low calcium intrapipette solution, as detailed in Methods. Representative amperometric traces for endophilin TKO, and endophilin TKO expressing endophilin 2 are shown in **Figure 3H**; the analyzed properties of the single spike are illustrated in **Figure 3I**. Significant differences were observed in the number of spikes/cell (**Figure 3J**) and in the single spike charge and amplitude (**Figure 3K-L**) - all these parameters were found to be decreased in endophilin TKO cells, while the foot properties and single spike rise time were not changed (**Figure 3M-Q**).

Given that exocytosis in chromaffin cells happens on the millisecond-to-second time scale, these data strongly suggest that endophilin has a direct role in exocytosis by controlling the sizes and fusion of releasable vesicle pools.

### LDCV vesicle morphology and number are not altered in the absence of endophilin-A

The decreased exocytosis in endophilin TKO chromaffin cells suggest that these cells might have less LDCV able to undergo exocytosis, or that the exocytic process itself is affected, or both. To distinguish between these possibilities, we performed comprehensive ultrastructural studies by combination of electron microscopy (EM) and 3D-confocal imaging.

Ultrastructural EM analyses on chromaffin cells in the adrenal gland were made as detailed in Methods. We noted that the overall size and morphology of endophilin TKO chromaffin cells was unaltered in comparison to controls (WT and littermate endophilin KOWTKO cells) (**Figure 4A)**, as well as the LDCV size (**Figure 4D**). Curiously, the average number of LDCVs per cell (**Figure 4B**), or per cell area (**Figure 4C**), were not significantly different between endophilin TKO, control littermates and WTs. This observation rules out the possibility that endophilin affects exocytosis by altering the number of LDCVs in the cell. Yet, while studying the distribution of LDCVs in endophilin TKO cells, we noticed that LDCVs were distributed further away from the plasma membrane in comparison to the controls (**Figure 4A** bottom panels, **Figure 4E-F’**), and less LDCVs were found within 10 nm from the plasma membrane (**Figure 4F**), suggesting a problem with vesicle recruitments to the release sites.

**Figure 4.**
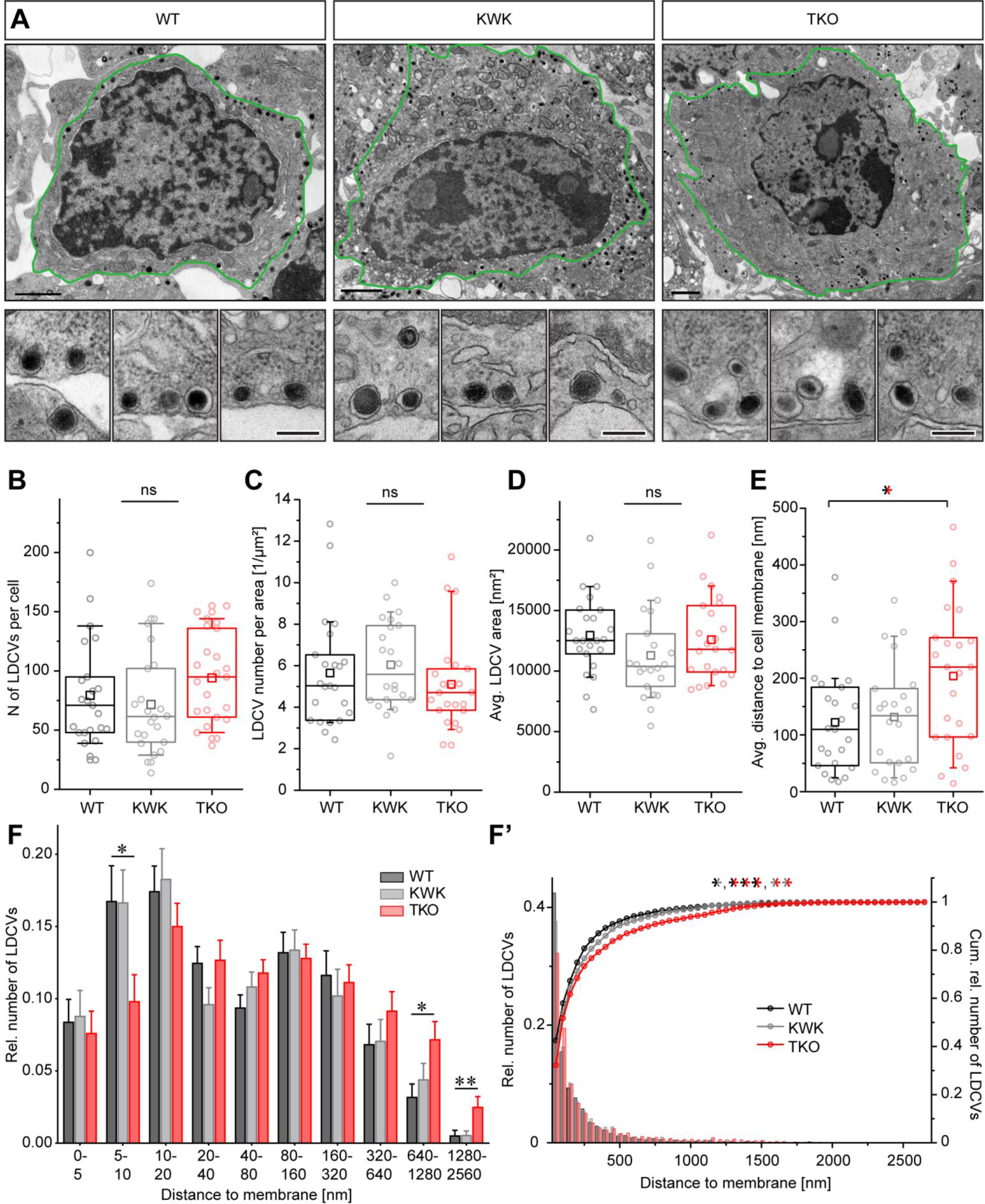
Number and size of LDCVs were not altered in endophilin TKO chromaffin cells, but less LDCVs was found in the plasma membrane proximity. (A) Example images of WT, endophilin KOWTKO (littermate control) and endophilin TKO chromaffin cells in the adrenal gland. Green line highlights one cell in the image. Note that the TKO cells is shown at lower magnification so the whole cell can be displayed. Panels below show higher magnification of LDCVs in the proximity of the plasma membrane (three examples are shown). Note that some LDCVs in endophilin TKO cells were not as close to the plasma membrane as in littermate and WT controls. Scale bar 100nm. (B-C) The total number of LDCVs per cell, and cell area, was unchanged between endophilin TKO and the control cells (one-way ANOVA). (D) The average LDCV area was unchanged between endophilin TKO and the control cells (one-way ANOVA). (E) The average distance of LDCVs from the plasma membrane was increased in cells without endophilins (one-way ANOVA followed by Tukey’s post-hoc test). (F) Distances between the LDCV membrane and the plasma membrane, after being normalized per cell. WT - black, endophilin KOWTKO - gray, endophilin TKO - red; at least 23 cells from 4 different animals and independent embeddings per group (Kolmogorov-Smirnov test with Bonferroni-correction). (F’) Relative frequency distribution and cumulative plots produced by binning all vesicles revealed altered distribution of LDCVs in endophilin TKO cells (Kolmogorov-Smirnov test with Bonferroni-correction).

The ultrastructural analysis was complemented by two independent approaches using confocal microscopy and Western blotting. First, fixed WT, endophilin KOWTKO and TKO chromaffin cells were immunostained against cargo marker CgA, and the whole cell (3D image acquired thorough z-stacks) was imaged by the Zeiss Airyscan system. Representative images are shown in **Figure 5A**. Quantification of LDCV number in the whole cell volume revealed similar number of LDCVs in endophilin TKO and controls (**Figure 5B**). Concordantly, the levels of CgA protein were not altered in endophilin TKO adrenal gland homogenates (**Suppl Fig S4A-B**).

**Figure 5.**
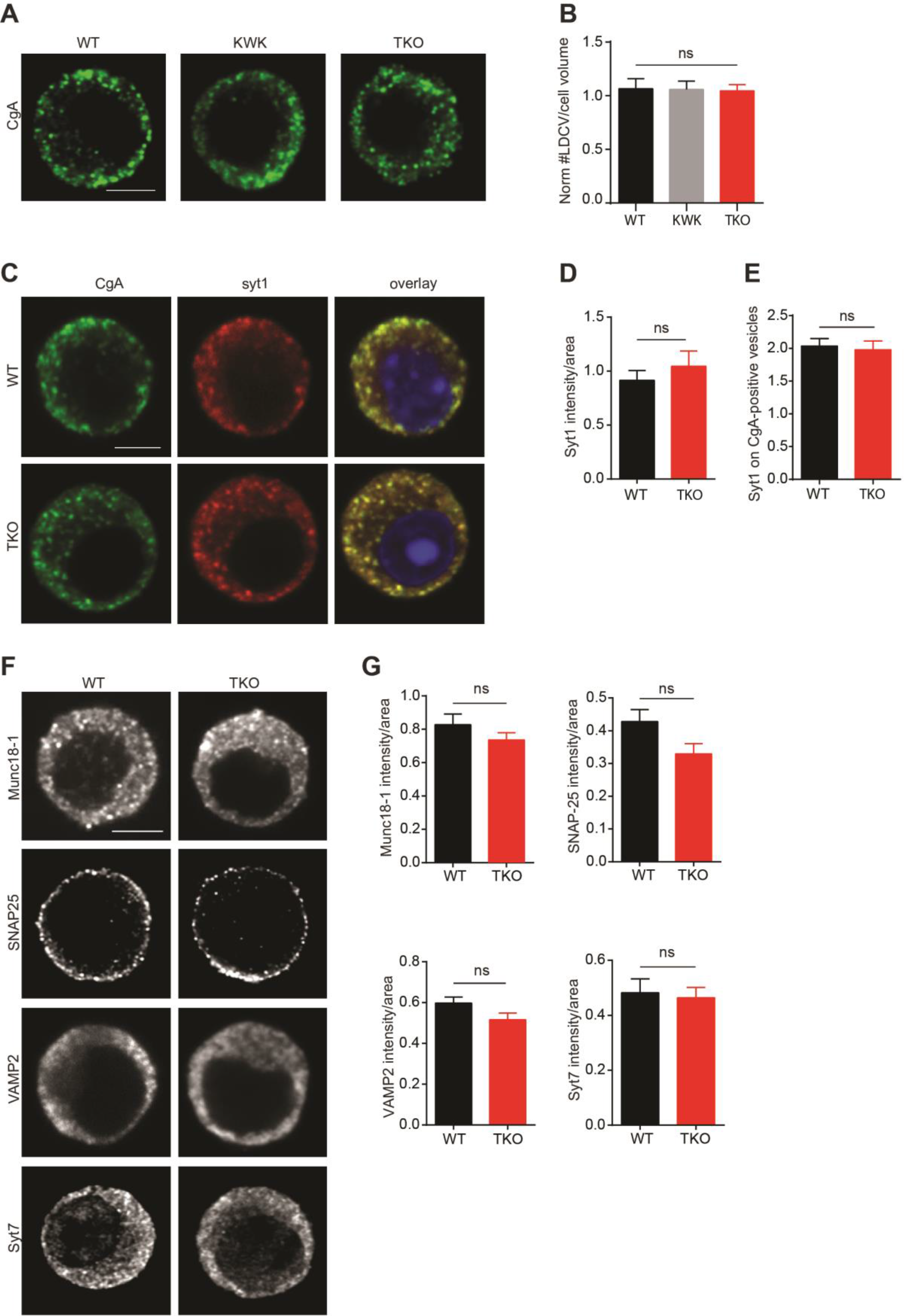
Exocytic machinery is unaltered in the chromaffin cells without endophilin. (A-B) Confocal image of endophilin TKO and control (WT and littermate endophilin KOWTKO) chromaffin cells stained with chromogranin-A (CgA; LDCV marker) revealed no difference in the number of CgA-positive vesicles measured in the whole volume of the cell (40 cells, p-value=0.96, one-way ANOVA with Tukey’s multiple comparison). An optical section through the equatorial plane of the cells is shown. (C) Images of endophilin TKO, littermate KOWTKO and WT chromaffin cells stained with anti-synaptotagmin-1 (Syt-1) and CgA antibodies. (D) Immunofluorescence levels revealed no change in synaptotagmin-1 levels (40 cells, p-value=0.45, unpaired t-test). (E) Quantification of the Syt-1 intensity on the CgA-positive vesicles revealed no significant difference between the genotypes (40 cells, p-value=0.76, unpaired t-test). (F) Representative images of endophilin TKO and WT chromaffin cell stained against Munc18-1, SNARE proteins SNAP25 and VAMP2 and synaptotagmin-7 shown no difference in the distribution of aforementioned proteins. (G) Quantification of Munc18-1 (30 cells, p-value=0.25, unpaired t-test), SNAP-25 (30 cells, p-value=0.07, unpaired t-test), VAMP-2 (40 cells, p-value=0.27, unpaired t-test) and Syt7 (30 cells, p-value=0.06, unpaired t-test) immunofluorescence showed no significant changes for any of these proteins.

Altogether, chromaffin cells without endophilin displayed minor alterations in the LDCV distribution, but the reduced exocytosis in these cells was not a result of the altered vesicle number, morphology or cargo (i.e. CgA) amount.

### Protein machinery involved in the exocytic process is not altered in chromaffin cells without endophilin-A

The reduced exocytosis (observed by fast electrophysiological measurements) in endophilin TKO cells occurs at the milliseconds-to-seconds time scale. Endophilins, being major endocytic proteins, could have an indirect effect on the LDCV composition and/or membrane and protein recycling processes in chromaffin cells. Despite the overall abundance of LDCVs and the unaltered morphology of individual LDCVs in endophilin TKO chromaffin cells, it is possible that some LDCVs may not be able to undergo exocytosis due to changes in accessory factors required for exocytosis. Specifically, SNAREs, Sec1/Munc-18 (SM)-proteins, Munc13s and other exocytic proteins may not be efficiently recycled from the previous rounds of exocytosis, or properly displayed on LDCVs in the absence of endophilin. Therefore, we checked the abundance and distribution of key exocytic proteins in endophilin TKO cells by immunocytochemistry (ICC) and Western blotting.

As an exemplary exocytic protein, we first analyzed the abundance and distribution of synaptotagmin-1 (Syt-1), a Ca^2+^-sensor important for LDCV exocytosis in chromaffin cells. We immunostained endophilin TKO, endophilin KOWTKO and WT cells for Syt-1, and noticed that neither distribution (**Figure 5C-D**), nor protein levels **Suppl Fig S4A-B**), of Syt-1 were altered in endophilin TKO cells. We next checked whether proper amounts of Syt-1 are present on LDCVs by quantifying the intensity of Syt-1 signal on the CgA-positive structures. The analysis showed no statistical difference of Syt-1 intensity on CgA-positive LDCVs between WT and endophilin TKO cells (**Figure 5E**).

We further inspected protein levels of Syt-1, as well as synaptophysin, several SNAREs (SNAP-25, syntaxin-1, synaptobrevin-2/VAMP2) and Munc-18-1 in adrenal gland homogenates by Western blotting, and found no significant difference between endophilin TKO cells and controls (littermate endophilin KOWTKO and WT samples were used as controls; **Suppl Fig S4 A-D**). Next, the ICC for a number of exocytic proteins, namely SNAP-25, synaptobrevin-2/VAMP-2, Munc18-1 and synaptotagmin-7 revealed no significant changes in the protein level and distribution in endophilin TKO chromaffin cells (**Figure 5F-G**). Taken together with the unchanged number of LDCVs, these data suggest that the reduced catecholamine release in the absence of endophilin is likely a consequence of endophilin’s direct action in exocytosis.

### Endocytic defects in the absence of endophilin-A in chromaffin cells

Endophilin TKO cells show an unaltered number of LDCVs and normal distribution and abundance of main exocytic proteins, yet, it is possible that altered endocytosis affects exocytosis in chromaffin cells. While exocytosis is well studied in this model system, endocytic modes are far from understood. Two temporally and mechanistically distinct forms of endocytosis have been reported: rapid endocytosis that depends on dynamin-1 and GTP, and slow endocytosis that involves dynamin-2 and clathrin (Smith and Neher, 1997; Elhamdani et al., 2006). We predominantly studied slow (likely clathrin-mediated) endocytosis since it is not known to which extent these cells undergo fast local recycling.

First, we examined the protein levels and distributions of main endocytic factors, namely clathrin heavy chain (HC), adaptor protein 2 (AP2), adaptor protein 180 (AP180), dynamins 1, 2 and 3 by ICC and Western blotting. Except for clathrin HC whose levels were mildly elevated in the absence of endophilin (by ICC only, **Figure 6A**), we detected no difference in the overall levels of aforementioned proteins, both by Western blotting of adrenal gland homogenates (**Figure 6B**) or by quantifying immunofluorescence in chromaffin cells (**Figure 6A**). In addition, the distribution of AP2 and dynamin seemed unaltered in endophilin TKO cells (**Figure 6A**).

**Figure 6.**
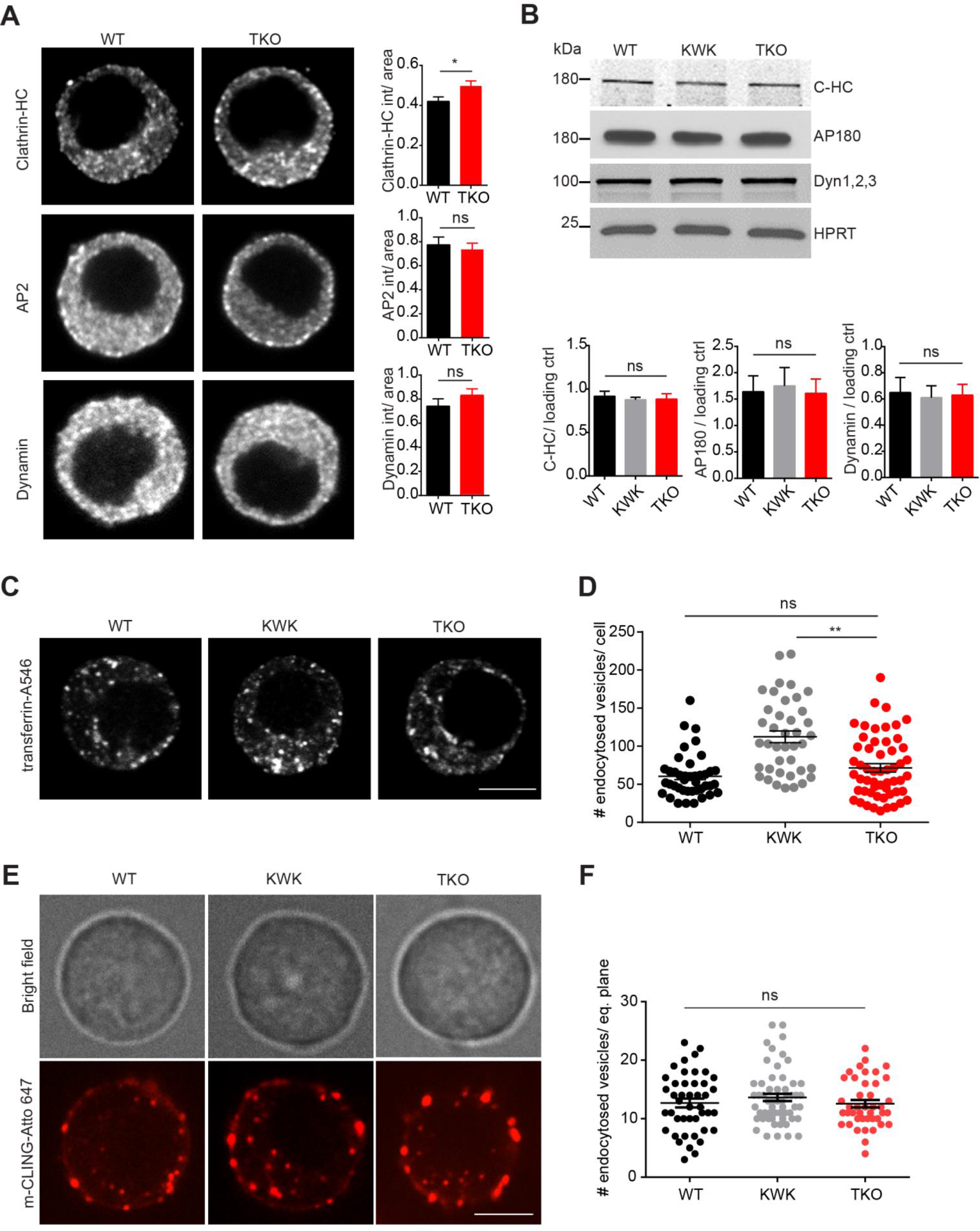
Endocytic defects in the absence of endophilin in chromaffin cells. (A) Immunofluorescence against clathrin heavy chain (HC), adaptor protein 2 (AP2) and dynamin-1 in WT and endophilin TKO cells. Fluorescence quantification revealed small but significant increase in clathrin HC intensities, whereas protein levels of AP2 and dynamin-1 were unaltered in endophilin TKO cells. (B) Proteins levels of main endocytic factors – clathrin, adaptor protein 180 (AP180) and dynamins 1-3 (inspected by Western blotting) were not altered in adrenal gland homogenates. Below: Quantification from 3-5 independent experiments. (C-D) Transferrin (conjugated with Alexa Fluor-546) uptake revealed a reduction in constitutive endocytosis in the endophilin TKO cells compared to the littermate control (p-value=0.001, one-way ANOVA, 40 cells; note that an uptake in the whole cell was analyzed). (E-F) mCling-Atto647 uptake (example cells shown in Movies 1, 3 and 4) by chromaffin cells of indicated genotypes showed no significant difference in the number of endocytosed vesicles (p-value=0.44, one-way ANOVA, 42 cells; note that an uptake in only one cell plane was analyzed). The specificity of mCling-Atto647 uptake was tested in the stimulated cells in the presence of Pitstop-2 inhibitor (Movie 2; Suppl data S5).

The slow endocytic recycling process in chromaffin cells was tested by three approaches: the uptake of transferrin-Alexa Fluor™ 546 (A546; clathrin-dependent) and uptake of mCling-Atto647 or recombinant cholera toxin subunit B (CT-B)-Alexa Fluor 594 (A594; clathrin-dependent and -independent). First, we looked at the 10 min-uptake of transferrin-A546 by analyzing 3D images (through z-stacks) of whole cell volume of endophilin TKO, endophilin KOWTKO and WT cells. The results were peculiar: while no significant difference between WT and endophilin TKO cells was observed (**Figure 6C-D**), endophilin TKO showed reduced amount of internalized transferrin-A594 when compared to endophilin KOWTKO cells (**Figure 6C-D)**.

We further inspected the uptake of the mCling-Atto647 that binds to the plasma membrane and whose internalization can be readily monitored for minutes (Revelo et al., 2016; **Suppl Fig S5A, Movie 1** shows stimulated chromaffin cells imaged for 8 min; for more information on the assay see Supplemental data). We first characterized the specificity of the mCling uptake in chromaffin cells by stimulating cells with high potassium in the presence of Pitstop-2 (clathrin coat formation inhibitor) and found that this inhibitor blocked the uptake of mCling efficiently (**Suppl Fig S5B, Movie 2 –** note that the cell surface increases upon stimulation since endocytosis was blocked). When the mCling-Atto594 was applied to chromaffin cells without endophilin, the number of internalized vesicles (mCling-positive structures) was comparable to controls (WT and endophilin KOWTKO cells; **Figure 6E**; **Movie 3** and **Movie 4**). Although a delay in kinetics in the first few minutes is a possibility in the TKO cells, the similar number of internalized vesicles was detected at 8 minutes (**Figure 6F**). A similar result was obtained with the uptake of CT-B-A594 (**Suppl Fig S5C-D**). Altogether, these data indicate that the lack of endophilin affects endocytosis in chromaffin cells only modestly and hence cannot account for the observed effect on exocytosis in this model system.

### Endophilin’s BAR-domain is not sufficient to mediate exocytic release from chromaffin cells

To get mechanistic insights on how endophilin regulates the exocytic process, we looked at the function of endophilin domains. Endophilins have two domains separated by a linker region: an N-terminal BAR-domain that senses and introduces membrane curvature and a C-terminal SH3-domain that mediates protein-protein interactions (e.g. with dynamin, synaptojanin-1, etc.). In nematodes, it has been shown that the endophilin’s BAR domain is necessary and sufficient to mediate the role of this protein in endocytosis (Bai et al., 2010), while in mammalian cells both domains were needed (Milosevic et al., 2011). We first tested whether endophilin’s BAR-domain alone (i.e. endophilin without SH3-domain) is sufficient to support exocytosis in chromaffin cells. We expressed endophilin 1 BAR-domain and endophilin 2 BAR-domain respectively (together with EGFP through the bicistronic IRES-expression system - the expression levels of all tested proteins were comparable; **Figure 7I**) in endophilin TKO cells, and performed capacitance and amperometry measurements as before. Interestingly, expression of endophilin 1 BAR-domain, as well as endophilin 2 BAR-domain, did not result in a rescue but rather in a further reduction of secretion from endophilin TKO cells, either during the first (**Figure 7A-D)** or the second stimulus **(Figure 7E-H**). The small responses revealed overall decrease in exocytosis, including both burst and the sustained component (first stimulus: **Figure 7B-D**; second stimulus **Figure 7F-H**). This dominant-negative effect reveals that the SH3-domain-mediated function is important for endophilin’s role in exocytosis, and that the full-length protein is needed to support exocytosis in chromaffin cells.

**Figure 7.**
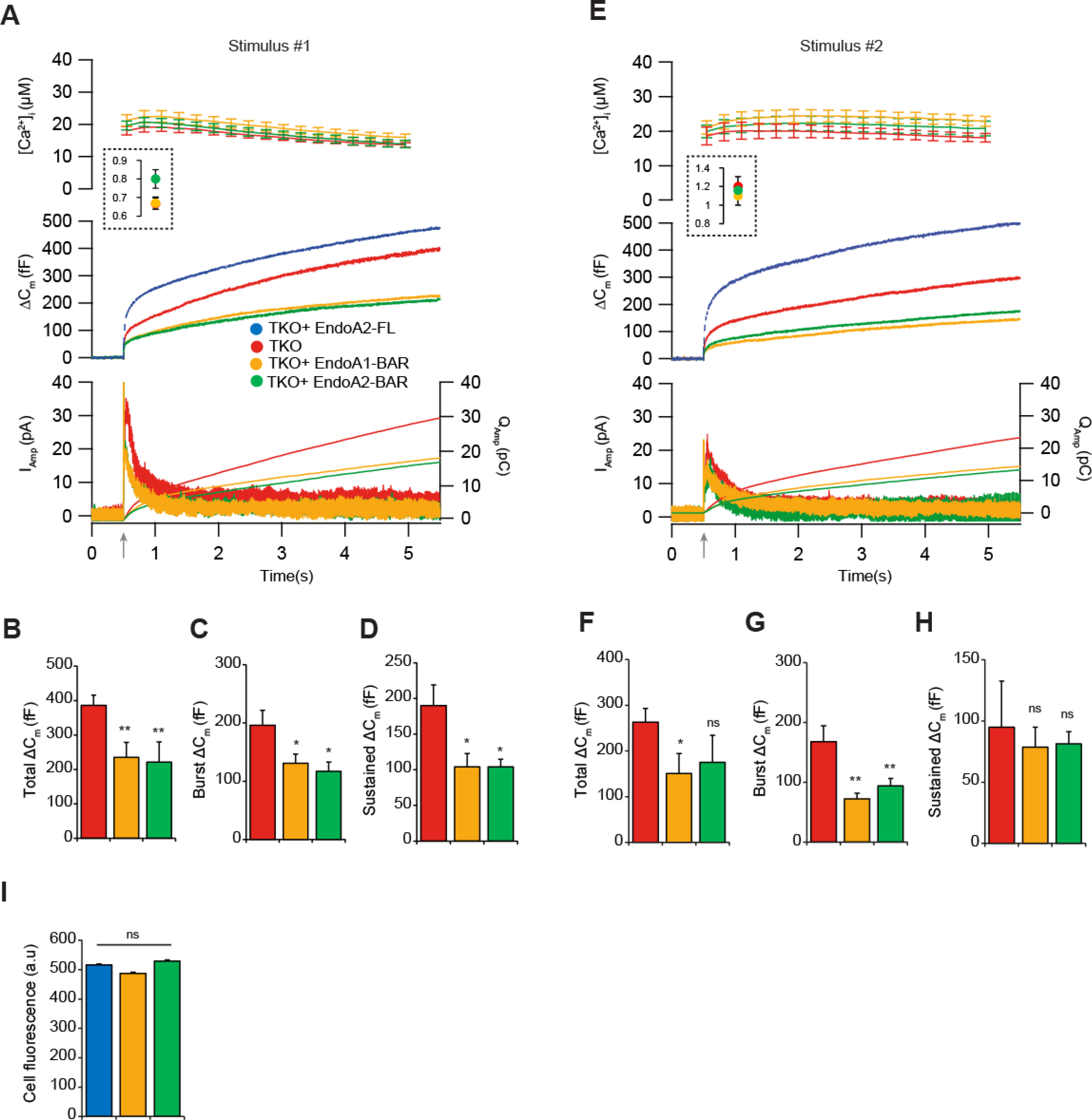
Endophilin 1 BAR and endophilin 2 BAR-domains are not sufficient to mediate exocytic release from chromaffin cells. (A) Exocytosis induced by calcium uncaging in endophilin TKO chromaffin cells compared to TKO cells expressing either endophilin 1 BAR or endophilin 2 BAR domain. Top: intracellular calcium level increase induced by flash photolysis at 0.5s (at arrow). The inset shows the pre-flash calcium levels. Middle: averaged traces of membrane capacitance upon Ca^2+^-induced exocytosis. Bottom: mean amperometric current (left axis) and cumulative charge (right axis). (C-H) Quantification of changes in capacitance revealed a further reduction in different phases of release - capacitance at 1s (burst) and 4.5s (total) in TKO cells expressing endophilin 1 BAR or endophilin 2 BAR domain (20 cells, one-way ANOVA with Tukey’s multiple comparison test). (I) Fluorescence intensities measured from cells expressing endophilin 2 (FL), endophilin 1 BAR or endophilin 2 BAR domain were comparable (48 cells).

### Endophilin’s role in exocytosis is mediated, at least in part, through its interaction with intersectin

Endophilin’s SH3-domain is known to mediate its interaction with several proteins, yet only two of them have been implicated in chromaffin cell exocytosis: dynamins (González-Jamett et al., 2010; Chan et al., 2010; Anantharam et al., 2011) and intersectins (Malacombe et al., 2006; Yu et al., 2009; Momboisse et al., 2010; Gerth et al., 2017). Distribution and levels of dynamins were not altered (**Figure 6A-B**), as detailed above. Curiously, while levels of intersectin-1, a membrane-associated protein that coordinates exocytic and endocytic vesicle traffic, were comparable (**Suppl Fig S6A-B;** both short and long isoform of intersecin-1 are shown), the distribution of intersectin-1 was altered in chromaffin cells lacking endophilins (**Figure 8A;** the line-intensity profile through the depicted cells is shown below the images). A detailed examination in resting chromaffin cells revealed that the fraction of intersectin-1 on the plasma membrane was higher in the TKO compared to the WT (**Figure 8B; Suppl Fig S6C-D**). Upon stimulation, more intersectin-1 was recruited to the plasma membrane in WT cells, whereas this redistribution did not happen in endophilin TKO cells (**Figure 8B; Suppl Fig S6C-D**). Similar observations were made with intersectin-2 (**Figure 8C-D; Suppl Fig S6E-F**). In sum, intersectin 1 and 2 are recruited to the plasma membrane (1) during stimulation and (2) in absence of endophilins.

**Figure 8.**
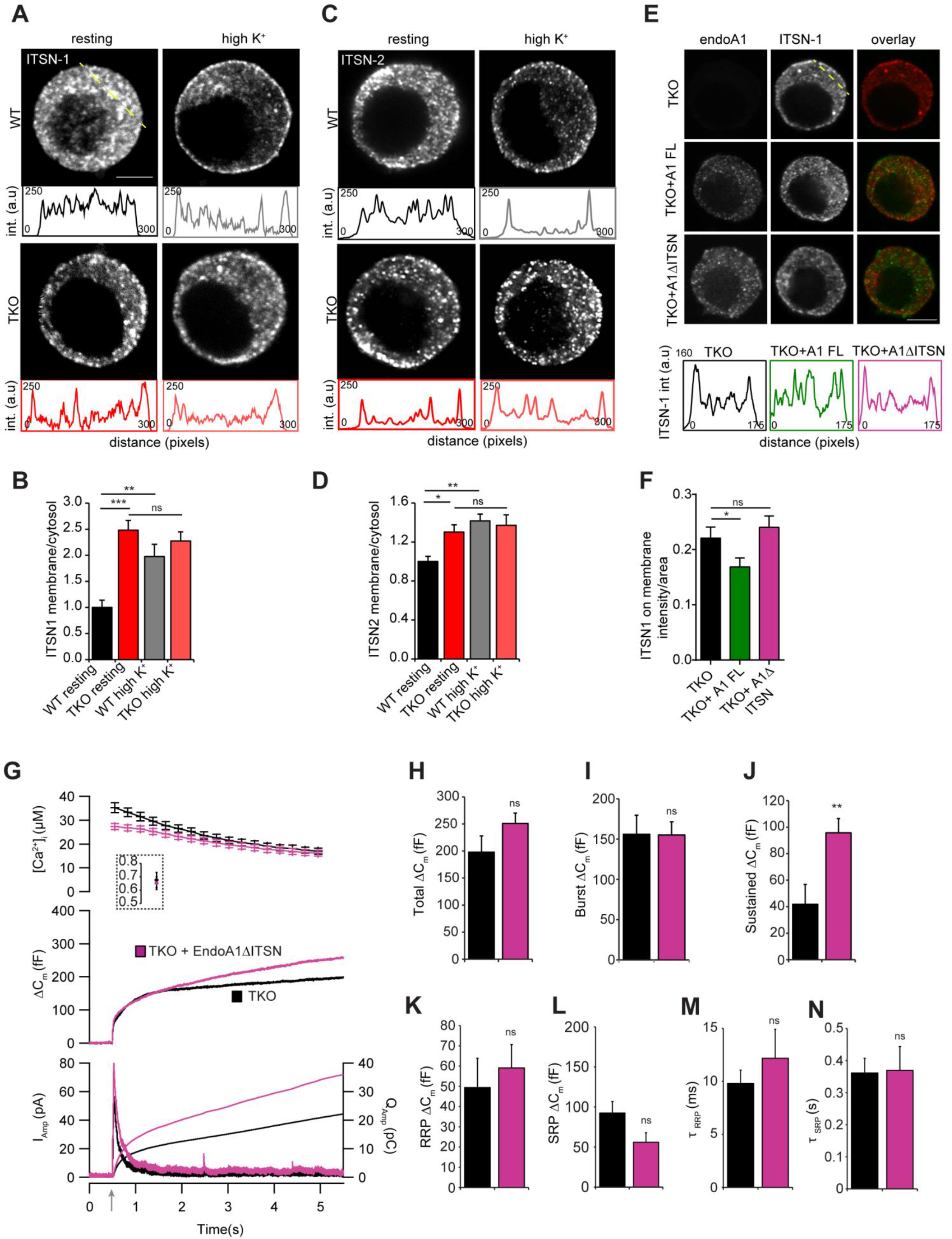
Endophilin’s role in exocytosis is mediated, at least in part, through its interaction with intersectin-1. (A-B) Distribution of intersectin (ITSN) 1 was altered in the endophilin TKO cells. (C-D) Distribution of ITSN-2 was altered in the endophilin TKO cells. Representative confocal images of chromaffin cells stained against ITSN-1 (A) and ITSN-2 (C) at resting and stimulated (depolarization by high K^+^) conditions. Intensity line profiles below indicate ITSN 1 and ITSN-2 intensities along the line (the approx. line position is marked in yellow in A). Quantification of ITSN-1 (B) and ITSN-2 (D) intensities in the cytosol vs. near the membrane (see Methods and Suppl Fig S6) revealed the altered distribution of ITSN-1 and ITSN-2 in endophilin TKO cells that did not further change upon stimulation (three independent experiments, p-value<0.0001, one-way ANOVA - multiple comparisons was done by Tukey’s multiple comparison test; note that WT resting vs. WT high K^+^, WT resting vs. TKO resting and TKO resting vs. TKO high K^+^ were compared). (E-F) ITSN-1 altered distribution in endophilin TKO cells was rescued by expression of endophilin 1, but not by expression of endophilin 1-ΔITSN (endophilin E329K+S366K mutant that does not bind ITSN-1). (G-N) Expression of endophilin 1 in endophilin TKO cells rescued exocytosis as inspected by combined capacitance and amperometry measurements (see Figure 3), but the same effect could not be achieved by expressing endophilin mutant that does not bind ITSN1 (E329K+S366K).

To check if this effect is specific, we attempted to rescue the intersectin-1 distribution by expressing either endophilin 1 or endophilin 1-E329K+S366K (mutant that does not bind intersectin; Pechstein et al., 2015) in endophilin TKO cells. Upon endophilin 1 expression in TKO cells, the distribution of intersectin-1 resembled the protein distribution in WT cells (as detected by immunostaining - **Figure 8E,** compare to **Figure 8A,** quantification in **Figure 8F**). Remarkably, the expression of mutant endophilin 1-E329K+S366K did not have the same effect, and intersectin-1 was still mislocalized (**Figure 8E-F**). These data suggest that endophilin-intersectin interaction is important for intersectin distribution in chromaffin cells and that it regulates intersectin’s access to the plasma membrane where the vesicle fusion happens.

Given that expression of endophilin 1 in endophilin TKO cells rescued exocytosis when inspected by combined fast capacitance and amperometry measurements (**Figure 3A-G**), we tested whether the same effect could be achieved by expressing the endophilin mutant that does not bind intersectin-1 (E329K+S366K, indicated as endoA1-ΔITSN). The expression of endoA1-ΔITSN through bicistronic lentiviral system was tested first (**Suppl Fig S6H**). Remarkably, endophilin 1-ΔITSN was not able to fully rescue exocytosis (**Figure 8G-I**), including the size and the time constants of vesicle pools (**Figure 8K-N**, note that **Figure 8J** depicts the sustained component that was at least partially rescued by endophilin 1-ΔITSN). Thus, endophilin’s role in exocytosis is mediated, at least in part, through its interaction with intersectin-1.

## DISCUSSION

The first reports on endophilin linked its function to endocytosis (Ringstad et al., 1997; de Heuvel et al., 1997; Ringstad et al., 1999; Verstreken et al, 2002; Schuske et al, 2003). Over three hundred papers in the past 20 years built on these initial findings and helped to unveil the mechanisms of endophilin action in several types of endocytosis, including clathrin-dependent (Ringstad et al., 1999; Verstreken et al, 2002; Schuske et al, 2003; Milosevic et al., 2011), clathrin-independent (Boucrot et al, 2015; Renard et al, 2015; Simunovic et al., 2017; Boucrot et al, 2018) and ultrafast endocytosis (Watanabe at al., 2018).

Our study shows that, in addition to its role in endocytosis, endophilin also plays a key role in the recruitment, priming and fusion of neurosecretory vesicles (endophilin 1 and 2 have overlapping functions in exocytosis). Endophilin is a peripheral protein with membrane-binding properties that appears to be present on at least some secretory vesicles: we detected its presence on SVs and LDCVs as well as a significant colocalization between LDCV marker CgA and endophilin 1 and endophilin 2, respectively. Upon stimulation, a fraction of endophilin translocates to the plasma membrane. Endophilin’s role in exocytosis is found to be mediated through its SH3-domain, most likely through its SH3-domain interactor intersectin, another membrane-associated protein that coordinates exocytosis and endocytosis and also translocates to the plasma membrane upon stimulation (see below). In the absence of endophilin, a significant fraction of intersectin is mislocalized to the plasma membrane, suggesting that endophilin acts as a repressor of intersectin by keeping intersectin away from the plasma membrane.

### Endophilin’s role in exocytosis is direct and endocytosis-independent

We consider that endophilin’s role in exocytosis is direct: The expression of either endophilin 1 or endophilin 2 alone was sufficient to rescue all exocytic defects seen in neurosecretory cells without endophilin. These data also show that two endophilins have a redundant role in exocytosis. In addition, the expression of endophilin 1 BAR domain, or endophilin 2 BAR domain, was not sufficient to produce a rescue of exocytosis in endophilin TKO cells. Next, endophilin that cannot bind intersectin-1 (E329K+S366K; Pechstein et al., 2015) was not able to fully rescue exocytosis. Lastly, none of the main exocytic proteins tested was found to be altered in the cells without endophilin.

The newly discovered role of endophilin in exocytosis appears to be independent of its well-established functions in endocytosis: the recycling/uptake of proteins (e.g. transferrin, cholera toxin subunit-B uptake) and membrane (fluorescently-labelled mCling reported in Revelo et al. (2016) was characterized through the Pitstop-2 application and stimulations, and found to be specific) were not majorly altered. We observed a mild decrease in the transferrin uptake efficiency between endophilin TKO and endophilin KOWTKO cells, yet no difference was detected between endophilin TKO and WT. This is a peculiar finding since it suggests that endophilin KOWTKO cells were more efficient in the transferrin uptake than WTs. Notably, the mCling uptake by TKO chromaffin cells was delayed in the first few minutes (Movie 1, 3, 4; although it was not found to be significantly altered at 8 minutes), so it is possible that the release site-clearance could be affected (Hua et al., 2013). Given that the LDCV generation/maturation steps take tens of minutes to hours, the short initial endocytic delays are likely not relevant for the recycling of proteins and generation of new LDCVs.

Despite the key exocytic (SNAREs, synaptotagmins and Muncs) and endocytic (AP2, AP180 and dynamin) machineries were not altered in endophilin TKO chromaffin cells, it was necessary to check whether all LDCVs were fusogenic (i.e. whether SNAREs and other exocytic proteins were efficiently recycled from previous rounds of exocytosis, and whether they were properly localized at the LDCVs). A systematic evaluation of each CgA-positive vesicle for synaptotagmin-1 presence and intensity revealed no major changes in the endophilin TKO chromaffin cells compared to the controls. In addition, we did not observe any overall differences in the levels and distributions of several additional vesicular proteins (e.g. VAMP2/synaptobrevin-2, synaptophysin, etc.).

### Endophilin’s role in exocytosis is mediated through its SH3-domain and intersectin interaction

We followed several leads that could explain endophilin’s role in exocytosis. The absence of endophilin’s SH3-domain reduced exocytosis even further than seen in endophilin TKO chromaffin cells, revealing a dominant negative effect of the BAR domain alone. Interestingly, out of all exocytic and endocytic (except clathrin) proteins inspected, only the distributions of intersectin-1 and intersectin-2 were altered. We reported previously that endophilin-intersectin-1 interaction is mediated through the SH3-domains of both proteins (Pechstein et al, 2015). We now found that an endophilin mutant that cannot bind intersectin-1 (E329K+S366K) was also not able to rescue the altered intersectin-1 distribution nor exocytic phenotype in endophilin TKO chromaffin cells. Both intersectin-1 and intersectin-2 were initially discovered as scaffold proteins involved in endocytosis (Yamabhai et al., 1998; Hussain et al., 1999; Simpson et al., 1999; Koh et al., 2004; Marie et al., 2004; Pechstein et al., 2010; Sakaba et al, 2013). Yet, further investigations showed that intersectins are also implicated in several other processes, including exocytosis (Malacombe et al., 2006; Yu et al., 2009; Momboisse et al., 2010; Sakaba et al, 2013; Gerth et al., 2017), thereby suggesting a role for intersectins in the coupling of exocytosis and endocytosis (Okamoto et al., 1999; Gubar et al., 2013; Gerth et al., 2017). Curiously, we observed that a fraction of intersectin-1 and intersectin-2 could be found on purified LDCVs and SVs (**Suppl Fig S6G**). Taken together with previous reports, our data suggest that endophilin regulates vesicle recruitment and exocytosis through its interaction with intersectin.

Both EM and immunofluorescent analyses of CgA-stained vesicles showed no changes in the number of secretory vesicles, thus, the reduction in exocytosis was not due to a reduced number of vesicles. However, curiously, the lack of endophilins had also altered the distribution of LDCVs near the plasma membrane. It is well known that the neuroendocrine cells (e.g. chromaffin cells) have a cortical F-actin barrier that controls the access of secretory vesicles to the plasma membrane (Trifaró et al., 1992). The polymerisation of the actin barrier seems primarily RhoA-dependent, while the *de novo* synthesis of actin is Cdc42-dependent (Malacombe et al., 2006; Momboisse et al., 2009). In addition, oligophrenin (a key RhoA inactivator) was shown to bind endophilin and assists its role in endocytosis (Nakano-Kobayashi et al., 2009; Nadif Kasri et al., 2011). We observed that, in the absence of endophilins, actin barrier looks depolymerized in the resting condition, while stimulation increased actin’s density (**Suppl Figure S7**) Thus, lack of endophilins led to a modulation of the actin barrier and the *de novo* synthesis. Given that actin is known to play a role in site clearance for vesicle docking and fusion (Miki et al., 2016), this perturbed actin dynamics may affect the availability of vesicles and their access to the plasma membrane, which in turn can explain the reduced exocytosis in the absence of endophilins. Therefore, the endophilin-intersectin interaction could also be a mechanism to control the actin barrier in chromaffin cells and subsequently the availability of vesicles for fusion. Once the actin barrier is depolymerized at the specific sites, secretory vesicles can be recruited to their site of release at the plasma membrane, possibly through an interaction between intersectin and the SNARE-protein SNAP-25 or SNAP-23 (Okamoto et al., 1999). Strikingly, chromaffin cells without intersectin-1 also showed reduced exocytosis (Yu et al., 2009), and intersectin was shown to regulate the replenishment of the fast-releasing synaptic vesicle pool in the calyx of Held synapse (Sakaba et al, 2013). We found that, without endophilin, intersectin mislocalized to the plasma membrane, LDCVs distribution was altered and exocytosis was diminished. In this context, endophilin may also be viewed as a regulator of intersectin in exocytosis.

### A putative model of endophilin’s role in exocytosis

This study, along with our previous results on endophilin and intersectin (Milosevic et al., 2011; Pechstein et al, 2015) and the reported interaction between intersectin and SNAP-25 and SNAP-23 (the SNARE proteins; Okamoto et al., 1999), as well as with the numerous intersectin studies (see above), suggests the following sequence of events (**Suppl Fig S8**). Endophilin is present at least on some neurosecretory vesicles and able to recruit intersectin. Endophilin and intersectin then act in tandem to stimulate the recruitment of those vesicles to the plasma membrane and their site of release, likely through intersectin’s role in the actin network modulation and its interaction with the SNARE proteins at the plasma membrane. Such recruitment may also be supported by the property of endophilins to bind and stabilize curved vesicular membrane via their BAR domain.

When the neurosecretory vesicle reaches the plasma membrane, the SNARE complex can be built between the vesicular VAMP2/synaptobrevin-2 and plasma membrane-resident SNAP25 and syntaxin-1, and endophilin and intersectin may stabilize this process. Once recruited to the proximity of the plasma membrane, we propose that endophilin and intersectin do not dissociate from the plasma membrane but take part in the endocytic process that follows the fusion of secretory vesicles. As such, endophilin and intersectin may act as a scaffold that couples vesicle fusion (exocytosis) and fission (endocytosis) events. This premise is supported by intersectin’s direct interaction with dynamin (Okamoto et al., 1999) and two studies in invertebrates: Bai et al. (2010) suggested that endophilin is delivered to endocytic zones by exocytosis in *C. elegans*, and Winther et al. (2013) showed that *D. melanogaster*’s Dap160/intersectin mutants lacking dynamin-binding do not properly accumulate dynamin in the periactive zone. Importantly, we show that intersectin mislocalizes to the chromaffin cell plasma membrane without endophilin. Since endophilin can regulate intersectin localization, it may be part of a check-point mechanism to ensure that intersectin acts only at the optimal time. This model is created primarily based on data obtained from neuroendocrine cells: it is possible that endophilin (and intersectin)’s roles in exocytosis diverge at the neuronal synapses where actin’s role may differ. Further studies are needed to address this question.

In conclusion, our study suggests that endocytosis of neurosecretory vesicles is dependent on endophilin 1 and endophilin 2, that were so far considered only as endocytic adaptors that form and stabilize membrane curvature. This novel function of endophilin is dependent on intersectin, and most likely actin.

## Supporting information

Movie 1. WT MCC with mCling

Movie 2. WT MCC with mCling and Pitstop2

Movie 3. Endophilin KWK-MCC-with mCling-100x

Movie 4. Endophilin TKO-MCC-with mCling-100x

## ACKNOWLEDGEMENTS

We thank the UMG animal facility, D. Schwitters, M. König and M. Costa for the excellent assistance and for help with genotyping, P. De Camilli (Yale University) and V. Haucke (FMP) for reagents and discussion, and A. Milosevic for drawing the model. This work was supported by Schram-Stiftung T287/25457 and Deutsche Forschungsgemeinschaft (Emmy Noether Young Investigator Award MI-1702/1) to IM, SySy fellowship to SG, the Lundbeck foundation (PSP, SH, JBS), ERC Starting Grant 337327 (NR), the Novo Nordisk Foundation (JBS) and the Independent Research Fund Denmark (SH, JBS).

## Author contribution

Conceptualization IM and JBS; Investigation and/or Analysis SG, VS, SH, MS, MG, NS, NR, and IM; Writing: SG, JBS and IM (all coauthors contributed to the final ms).

Authors declare no competing financial interests.

## STAR METHODS

### CONTACT FOR REAGENT AND RESOURCE SHARING

Further information and requests for resources and reagents should be directed to and will be fulfilled by the Lead Contact, Ira Milosevic.

### EXPERIMENTAL MODEL

All animal-related procedures were performed according to the European guidelines for animal welfare (2010/63/EU), with the explicit permission from the Niedersächsisches Landesamt für Verbraucherschutz und Lebensmittelsicherheit (LAVES), registration 14/1701. Animals were housed and bred in the Zentrale Tierexperimentelle Einrichtung (ZTE) Göttingen with *ad libitum* access to water and food. Mice were kept in groups of 1-5 animals on a 12h light/12h dark cycle in the individually ventilated cages. Endophilin mutant mice were originally generated and described in (Milosevic et al., 2011), and are accessible from the Jackson Laboratory (strain 021573 - B6;129-Sh3gl2^tm1Pdc^/J; strain 021574 - B6;129-Sh3gl1^tm1Pdc^/J; 021575 - B6;129-Sh3gl3^tm1.1Itl^/J). WT (C57BL/6J) mice used as additional control were obtained from endophilin A1^+/−^A2^+/−^A3^+/−^ mice breeding, or were purchased from ZTE. Both male and female P0 pups were used to prepare the culture of adrenal chromaffin cells as follows: Adrenal glands were isolated from P0 mice (endophilin TKO, endophilin KOWTKO or WT-C57/BL6J) and placed in Locke’s solution (154 mM NaCl, 5.6 mM KCl, 0.85 mM NaH_2_PO_4_, 2.15 mM Na_2_HPO_4_, and 10 mM glucose, pH 7.25), and cleaned to remove connective tissue. The glands were digested in 200 μl of enzyme solution at 37 °C for 30-35 minutes (enzyme solution: DMEM medium with high glucose (Gibco) supplemented with 25 mg L-cysteine, 125 μl 1 M CaCl_2_, 1.25 ml 50 mM EDTA pH 7.4, and 22.5 units/ml papain). The enzyme was inactivated by adding 200 μL inactivating solution for 5 minutes at 37 °C (inactivating solution: 112.5 ml DMEM medium with high glucose supplemented with 12.5 ml heat-inactivated fetal calf serum (Life Technologies), 312.5 mg albumin fraction-V, 312.5 mg trypsin inhibitor). This solution was replaced by 180 μL enriched DMEM medium and the glands were triturated (5-6 times) using a 200 μL pipette (enriched DMEM medium: 6.7 g DMEM powder with high glucose (Gibco) was dissolved in 496.5 ml double-distilled water supplemented with 0.55g NaHCO_3_, 1 ml penicillin/streptomycin (Life Technologies) and 2.5 ml insulin-transferrin-selenium-X (Thermo Fischer Scientific). The cells were plated on cleaned and sterilized glass coverslips and allowed to settle for 30-45 minutes before adding 1 mL enriched DMEM medium. The cells were maintained at 37 °C and 8 % CO_2_ and used within 3-4 days for the experiments.

## METHOD DETAILS

### Cloning and virus production

For the rescue experiments, full length endophilin 1, and endophilin 2, were cloned into lentivirus (LV) expression vector FUGW (a gift of Oliver Schlüter, European Neuroscience Institute Göttingen, Germany) containing an IRES followed by an enhanced green fluorescent protein (EGFP) to allow simultaneous yet independent expression of both proteins. Endophilin 1, and endophilin 2, were first amplified by a PCR reaction (original plasmids were described in Milosevic et al., 2011) and then inserted into the FUGW vector using *XbaI* and *BamHI* restriction enzymes. Similarly, endophilin 1-BAR and endophilin 2-BAR constructs (BAR domain and the linker sequence) were cloned by amplifying and inserting the endophilin 1-BAR and 2-BAR sequences into FUGW vector using *XbaI* and *BamHI* restriction enzymes. Endophilin 1-ΔITSN (endophilin 1 E329K+S366K - mutant that cannot bind intersectin-1; Pechstein et al, 2015) was first generated by QuikChange II Site-Directed Mutagenesis Kit (Agilent) and subsequently inserted into the FUGW vector using *XbaI* and *BamHI* restriction enzymes. All constructs were verified by sequencing and control restriction digestion. Lentiviral particles were generated as follows: 293FT cells were plated at 1×10^7^ per ø10cm dish. The cells were transfected with lentivirus transfer plasmid as detailed above (3^rd^ generation lentivirus system) along with envelop and packaging plasmids using Lipofectamine-2000 and following the manufacturer’s protocol (Invitrogen). The cells were maintained in the S2 bio-safety laboratory henceforth and the medium was exchanged 14h post-transfection. The medium containing lentivirus suspension was collected, centrifuged at 3,000 RPM for 15 minutes at 4°C to remove cell debris. Further, virus was concentrated using Amicon (100K, UFC910096) at 4,000 RPM for 20 minutes at 4°C. The concentrated particles were diluted up to 2mL of Tris-buffer saline (TBS; pH 7.4); aliquots were frozen in cryo-tubes in liquid nitrogen and stored in −80°C until being used. The efficiency of the lentivirus was tested by Western blot and imaging of a fluorescent reporter. The virus particles were added 6-8 hours after cell plating, and the cells were used 60-72 hours post infection.

### Purification of synaptic vesicles and LDCVs

Synaptic vesicles were purified as recently published (Farsi et al., 2018). LDCVs were purified according to Park et al. (2012) with minor modifications. Specifically, the medulla was isolated from two bovine adrenal glands obtained from a recently slaughtered animal, was minced in 300 mM sucrose buffer (300 mM sucrose, 20 mM HEPES, pH 7.35, 200 μM PMSF) and then homogenized using a cooled teflon-glass homogeniser (Dounce homogenizer, Wheaton) at 1,000 rpm. After centrifugation at 1,000 g for 15 min at 4°C, the pellet was discarded and the supernatant was subjected to another centrifugation at 12,000 g, 15 min, 4°C. This step was repeated once more, and the resulting pellet was resuspended in 300 mM sucrose buffer and loaded on top of a continuous sucrose gradient (from 300 mM to 2.0 M) to remove contaminants. LDCVs were collected from the pellet after centrifugation at 27,000 rpm for 60 min (Beckman SW 41 Ti rotor) and re-suspended in 120 mM K-glutamate, 20 mM K-acetate, 20 mM HEPES, pH 7.4. Protein concentration of SV and CCV samples was determined using Pierce BCA Protein Assay Kit.

### Electrophysiological measurements

The mouse chromaffin cells were maintained in extracellular solution (145 mM NaCl, 2.8 mM KCl, 2 mM CaCl_2_, 1 mM MgCl_2_, 10 mM HEPES, and 2 mg/ml D-glucose, pH 7.2, 305 mOsM/kg) during the electrophysiological recordings. Capacitance and amperometric measurements were performed in parallel on a Zeiss Observer.D1 equipped with Polychrome V monochromator (Till Photonics), an EPC-10 amplifier (HEKA Elektronik) for patch-clamp capacitance measurements and an EPC-9 (HEKA Elektronik) for amperometry. Catecholamine release was triggered by UV-flash photolysis (using a JML-C2, Rapp Optoelektronik) of a caged calcium compound, nitrophenyl-EGTA, which was transferred into the cell through a patch pipette. The intracellular calcium was monitored, after the setup calibration was done by infusion of 8 solutions of known calcium concentrations into chromaffin cells, by ratiometric measurement of two calcium dyes with different calcium binding affinity, Fura4F and Furaptra (ThermoFischer Scientific). The excitation light (Polychrome V) was alternated between 350 nm and 380 nm to perform the ratiometric measurements of [Ca^2+^]_i_. The emitted fluorescence was detected with a photodiode and sampled using Pulse software (HEKA). The same software was used to control the voltage in the pipette and perform capacitance measurements. The intracellular solution contained (in mM): 100 Cs-glutamate, 8 NaCl, 4 CaCl_2_, 32 Cs-HEPES, 2 Mg-ATP, 0.3 GTP, 5 NPE, 0.4 Fura4F and 0.4 Furaptra (L-ascorbic acid was added to prevent flash-induced damage to Fura dyes), pH 7.2 (osmolarity adjusted to ∼295 mOsm). Amperometric recordings were done using ø5 μm carbon fibers (Thornel P-650/42, Cytec) insulated by the polyethylene method (Bruns, 2004) and EPC-7 (HEKA). Cells were perfused with intracellular solution for single amperometry spikes, consisting of 70 mM Cs-glutamate, 8 mM NaCl, 4 mM CaCl_2_, 22.5 mM Cs-HEPES, 2 mM Mg-ATP, 0.3 mM Na-GTP, 37 mM Ca2^+^ DPTA, 0.32 mM Fura-4F and 0.48 mM Furaptra pH 7.2 (osmolarity adjusted to ∼300 mOsm). Fibers were clamped to 700 mV. Currents were acquired at 25 kHz and filtered off-line using a Gaussian filter with a cutoff set at 1 kHz.

### Immunocytochemistry

Chromaffin cells were cultured for up to 72h on poly-L-lysine coated coverslips in a 12-well plate (Sarstedt) for immunocytochemistry experiments. The cells were fixed in freshly prepared 4% paraformaldehyde (PFA) in phosphate buffered saline (PBS; Sigma) and neutralized afterwards for 20 minutes with 50mM NH_4_Cl in PBS. Blocking was performed with the blocking buffer containing 3% bovine serum albumin (BSA; Sigma), 0.2% cold-fish gelantine (Sigma) and 1% goat serum (Gibco) for 1 hour, and cells were permeabilized in 0.1% Triton X-100 (Sigma) in the blocking buffer for 10 minutes. After a brief washing step, the cells were stained with primary antibody overnight at 4°C followed by washing and secondary antibody staining in the dark at room temperature for 1h (the antibodies are listed in Key Resources Table). The washing procedure was repeated following 2 min incubation with 4’,6-diamidino-2-phenylindole (DAPI 1:5,000 in PBS; Sigma) to stain the nucleus of the chromaffin cells. Finally, the coverslips were mounted in Mowiol 4-88^®^ mounting medium (Sigma). Images were acquired using Zeiss LSM 710 laser scanning confocal microscope or Zeiss LSM 800 Airyscan confocal microscope (63x objective, numerical aperture 1.4).

For chromaffin cell stimulation, the cells were washed carefully in pre-warmed Locke’s solution before incubation in extracellular (control condition) or high K^+^ solution (88 mM NaCl, 59 mM KCl, 2 mM CaCl_2_, 1 mM MgCl_2_, 10 mM HEPES, and 2 mg/ml D-glucose, pH 7.20, 300 mOsM/kg) for 3 minutes at RT. The cells were placed on ice immediately, fixed in 4% PFA (freshly prepared) for 10 minutes on ice followed by 20 minutes at RT. Immunostainings and image acquisition were performed as described above.

### Electron microscopy on adrenal chromaffin cells

Adrenal glands from endophilin TKO, endophilin KOWTKO and WT P0 mice were isolated and subsequently fixed in 4% PFA + 0,5% glutaraldehyde (GA; Sigma) in 0.1M PBS, pH 7.2 for 1 hour on ice, and afterwards in 2% GA in 0.1M sodium cacodylate buffer (pH 7.2; Sigma) overnight at 4°C. The next day, the adrenal glands were washed 3 times for 10 minutes in 0.1M sodium cacodylate buffer. Post-fixation was done on ice for 1 hour in 1% (v/v) OsO_4_ in cacodylate buffer, followed by further washing steps (2×10 min cacodylate buffer, 3×5 min water). En-bloc staining of adrenal glands was performed using 1% (v/v) uranyl acetate (Sigma) in water for 1h on ice. Subsequent to three brief washing steps in water, adrenal glands were dehydrated in an ascending ethanol series (5 min 30%, 5 min 50%, 10 min 70%, 2×10 min 95%, 3×12 min 99,9% ethanol) and infiltrated in Epon resin (50% ethanol + 50% epon for 30 min and for 90 min, 100% epon for ~20h) at room temperature. The samples were placed in the embedding molds and polymerized for 48h at 70°C. Ultrathin sections (65 nm) were cut using a Leica UC6 ultra microtome, placed on formvar-coated copper grids, and stained in uranyl acetate and lead citrate (using the Reynold’s method). Images were acquired using a JEOL JEM-1011 transmission electron microscope equipped with a Gatan Orius 1200A camera at 6000x magnification, on average 5 images/cell were acquired.

### Endocytosis assays on chromaffin cells

Transferrin (Tf) was conjugated to Alexa Fluor™ 546 (5 μg/mL; Invitrogen, Cat.# T23364), and the uptake assay in adrenal chromaffin cells was performed as described in Chen *et al*. (1998). All images were captured under the same acquisition settings using a Zeiss 810 Airyscan confocal microscope. Analyses of Tf-A546 data (z-stack of the whole cells) was performed by Imaris (Bitplane) using the Spot module and statistics was performed by one-way ANOVA. Non-toxic recombinant cholera toxin subunit-B (CT-B) was conjugated to Alexa Fluor™ 594 (Thermo Fischer, Cat.# C22842) and the uptake assay was adapted to adrenal chomaffin cells using the protocol from Kirkham *et al*. (2005). In short, uptake of 2 g/mL CT-B-A594 was carried out at 37°C in serum-free medium (Gibco) for 8 min (note that CT-B-A594 attaches to cells by binding to ganglioside GM_1_), Cells were washed 4 × 30s with the extracellular buffer to remove CT-B-A594 cell surface labeling, fixed in 4% PFA for 20 min, and imaged with a Zeiss 810 confocal microscope. Images were captured through the equatorial plane of each cell (one plane only) using the same acquisition settings, and were analyzed in ImageJ (a background-threshold was applied and every fluorescent cluster greater than six pixels was counted).

Live imaging of chromaffin cells labeled with mCling-Atto647N (Synaptic Systems) was done using the spinning disk confocal microscope with temperature control unit (kept at 37°C) and custom-built imaging chamber. Cells were maintained in extracellular solution in imaging chamber and stained with 0.5 nmol/ml mCling-Atto647N for 1 minute. The solution was exchanged and cells were washed rapidly (few seconds) before image acquisition. Images were captured up to 8 minutes after addition of mCling-Atto647N through the middle of each cell (using the same acquisition settings), and quantified using ImageJ as detailed above for CT-B-A594. Statistics was performed by one-way ANOVA.

## QUANTIFICATION AND STATISTICAL ANALYSIS

Endophilin-CgA colocalization analysis in chromaffin cells was performed as follows: colocalization was evaluated using object-based overlap and JACoP plugin in ImageJ since endophilin signals had a cytosolic component. Specifically, a region-of-interest (ROI) was chosen so it did not contain nucleus, and the images were segmented into objects and background (bright fluorescent objects were segmented from the image) before Pearson’s correlation coefficient was calculated. The same analysis was performed with the 90 rotated image, and this random value was subtracted for each cell. In addition, the complementary manual colocalization analysis was performed: here, circles were superimposed on bright fluorescent spots in the CgA channel and transferred to identical image locations in the endophilin channel. If the fluorescence intensity maximum in the endophilin channel was located in the same circle and the morphology of the signal resembled that of the CgA signal, the circle was rated as positive (colocalized). If this was not a case, it was rated as negative (not colocalized). To correct for random colocalization of two abundant signals, circles were also transferred to a 90° rotated image of the endophilin channel. A minimum of 10 images from 3-5 experiments were analyzed for each genotype/condition. Two approaches (semi-automatic ImageJ-based and manual) gave similar result.

The kinetic analysis of the capacitance measurements was performed by fitting individual capacitance traces with a triple-exponential function using a custom-written macro in IGOR Pro, as described in Milosevic et al., 2005. The amplitudes and time constants of the two faster exponentials define the size and release kinetics of the slowly releasable pool (SRP) and the readily releasable pool (RRP), respectively. Filtering, spike detection and analysis of aperometric spikes were performed by a custom-written macro (Mosharov and Sulzer, 2005) in IGOR Pro (Wave Metrics). Data are represented as mean±SEM, and unpaired t-test or nonparametric Kruskal-Wallis test with Dunn’s multiple comparison test were used to test statistical difference, which is indicated by *p<0.05, **p<0.01, and ***p<0.001.

The ultrastructural analysis by EM: At least 23 cells from 4 different animals and independent embeddings per group were analyzed using the IMOD (bio3d.colorado.edu/imod) and ImageJ software. The area of chromaffin cells was defined as area within the plasma membrane, thereby excluding the nucleus area. The area of LDCVs was directly measured using the area selection tool in ImageJ, and distances were measured between LDCV membrane and plasma membrane in IMOD. The statistics is done by one-way ANOVA (Figure 4B-D) followed by Tukey’s post-hoc test (Figure 4E) and Kolmogorov-Smirnov test (Figure 4F-F’; to ensure that the detected differences were not artefacts created by multiple comparisons, we applied Bonferroni-correction).

For quantification of CgA-positive vesicles (Figure 5A-B) and Syt-1 intensity on CgA-positive vesicles (Figure 5C and 5E), three-dimensional surface reconstruction was carried out with the “Cell with organelles” module of Imaris software, version 8.0.2 (Bitplane).

For quantification of intensity of proteins detected by immunofluorescence (Figure 5D, Figure G and Figure 6A), the mean fluorescence intensity was measured in the cell cytoplasm, excluding the nucleus, using ImageJ and normalized to the measured area. The data are represented as mean±SEM and statistics was performed by unpaired t-test.

To characterize the distribution of intersectins (Figure 8A-F; Suppl Fig S6D-G), we capitalized on the round-shape of adrenal chromaffin cells, and defined a ROI-based analysis approach in ImageJ software. We defined two concentric ROIs: the outer circular ROI around the whole cell and the inner circular ROI (to measure the intensity of the cytosol). The inner circular ROI was defined to be 40 pixels less than the radius of the outer ROI. The intensities of two ROIs were then measured, and the inner circular ROI was subtracted from the outer circular ROI to calculate the near-membrane intensity. The ratio of intensities between membrane and cytosol was plotted. The data are represented as mean±SEM from three independnet experiemnts with at least 45 cells per group and the statistics was performed by one-way ANOVA.

