## Supplementary figures and images for "Endophilin-A controls recruitment, priming and fusion of neurosecretory vesicles"

### Supplemental figures

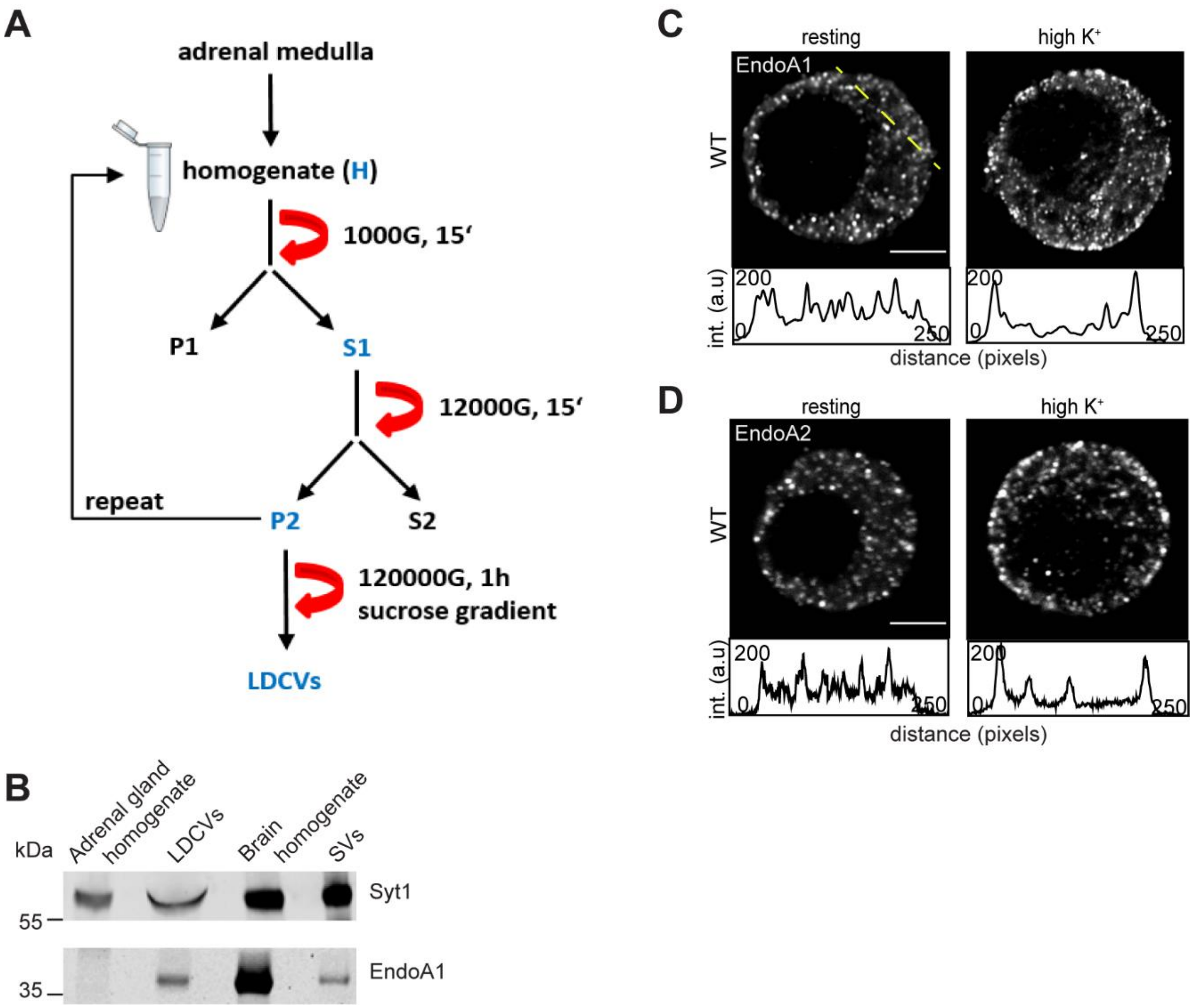

**Suppl. Fig. S2.**

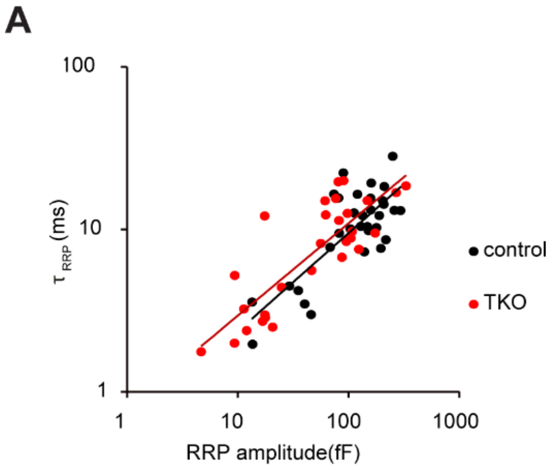

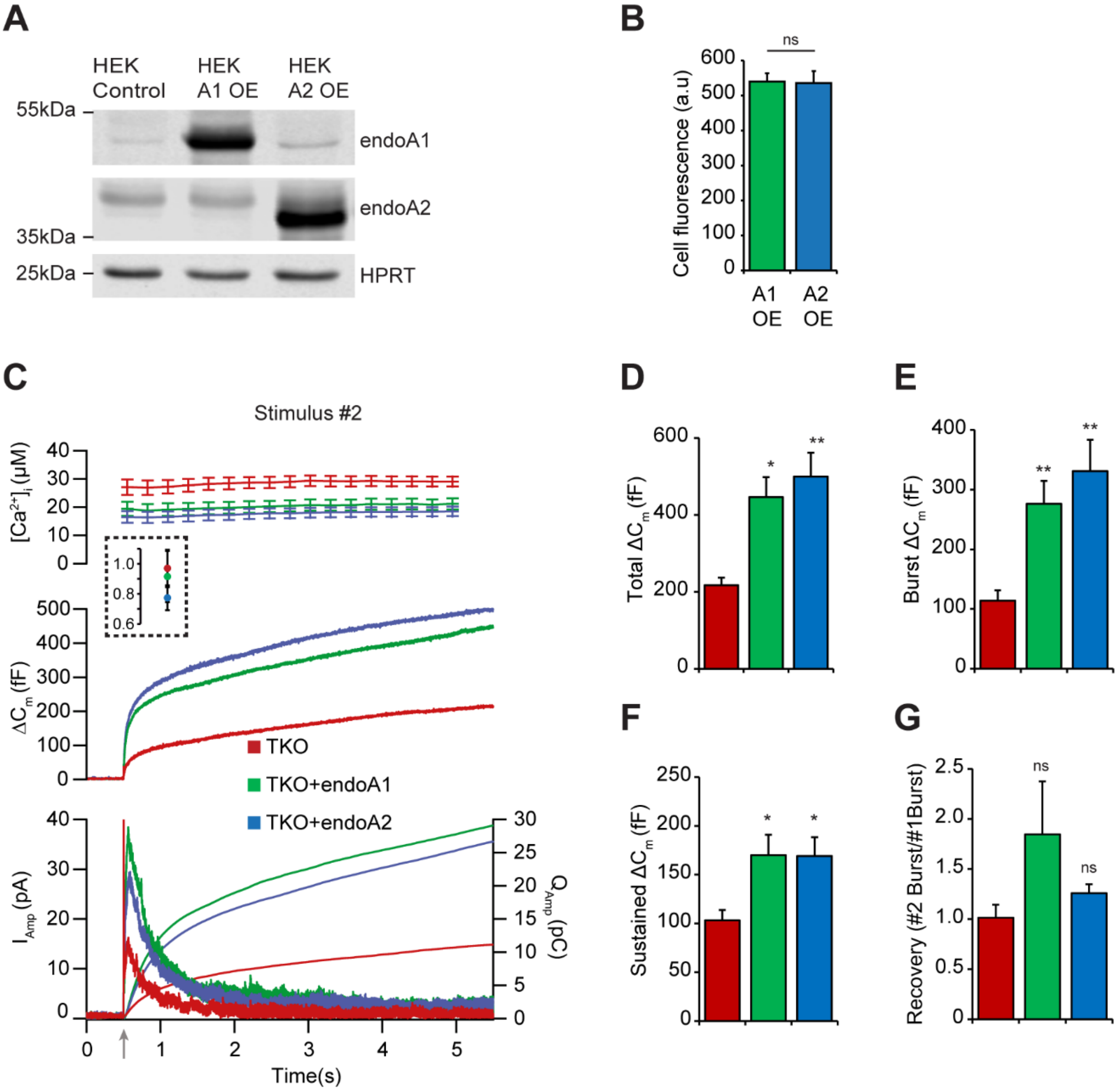

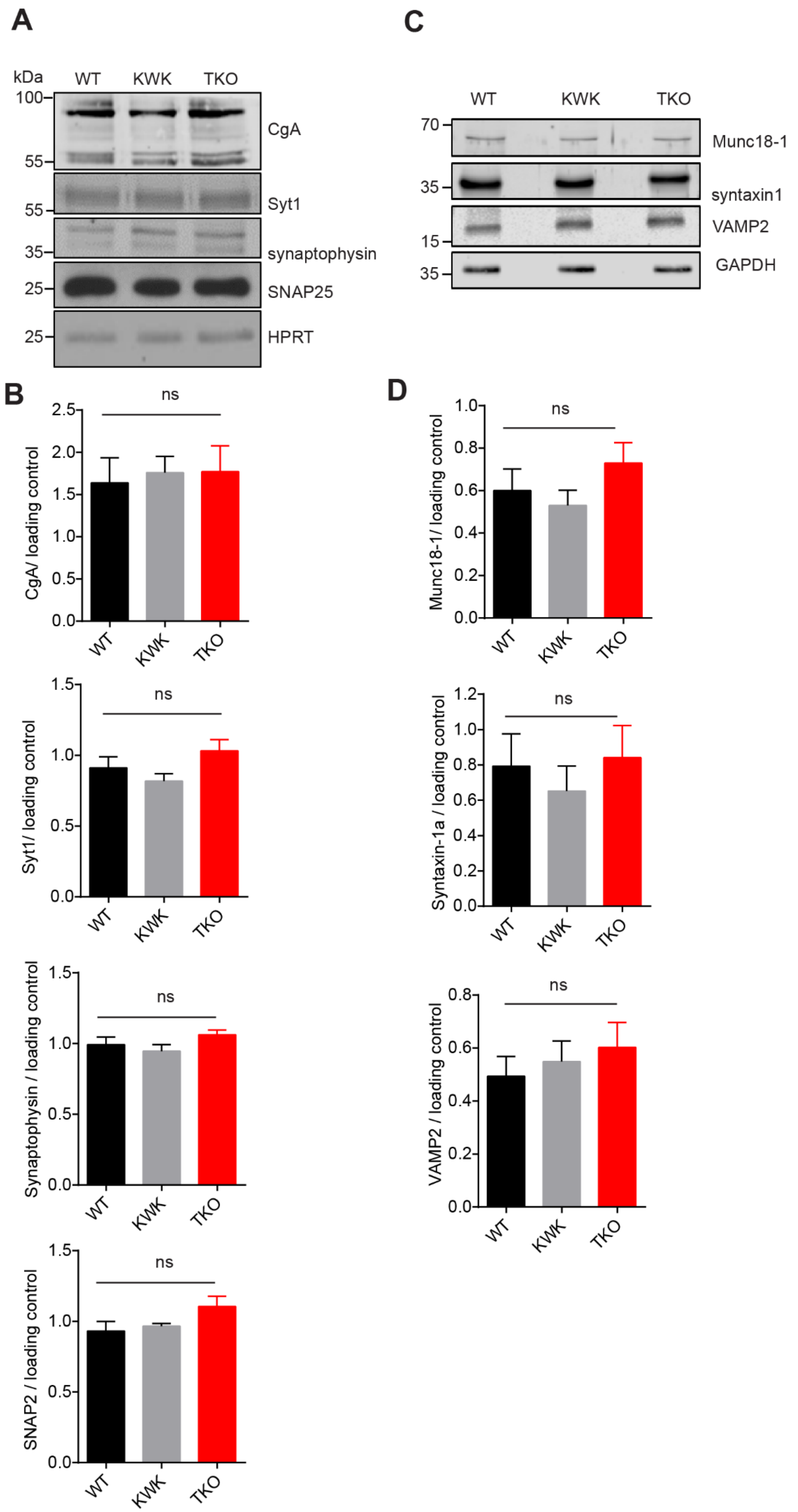

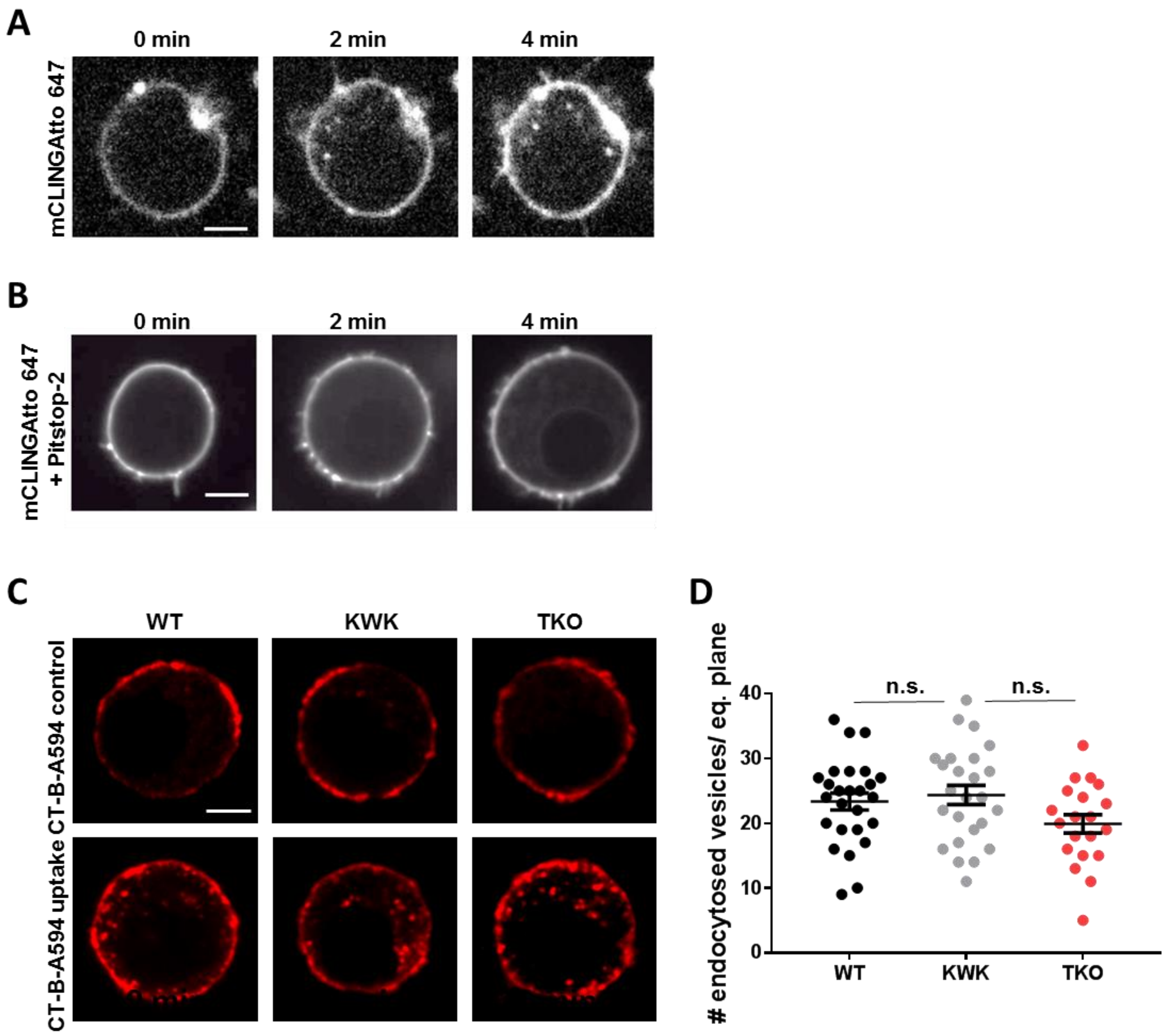

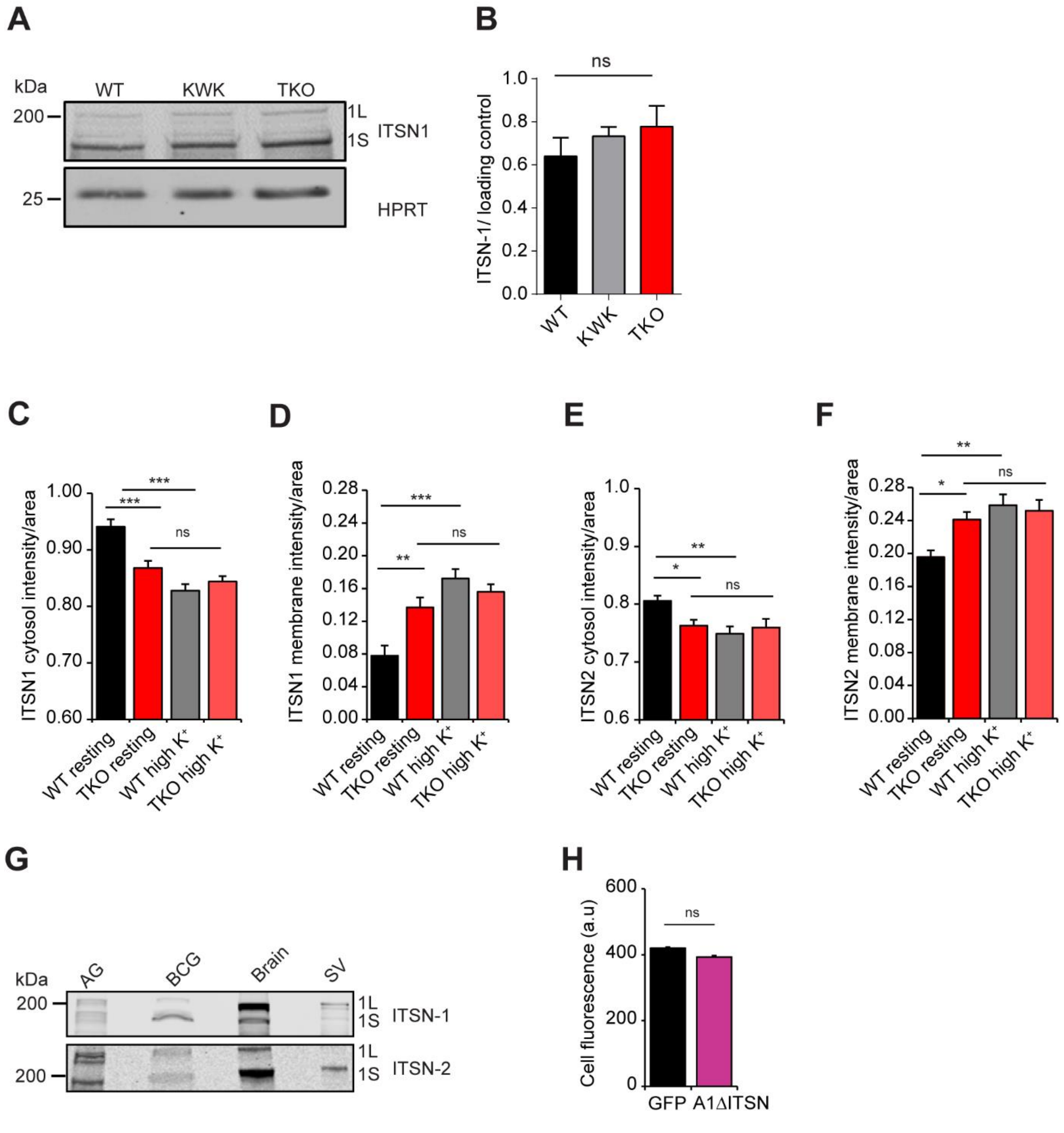

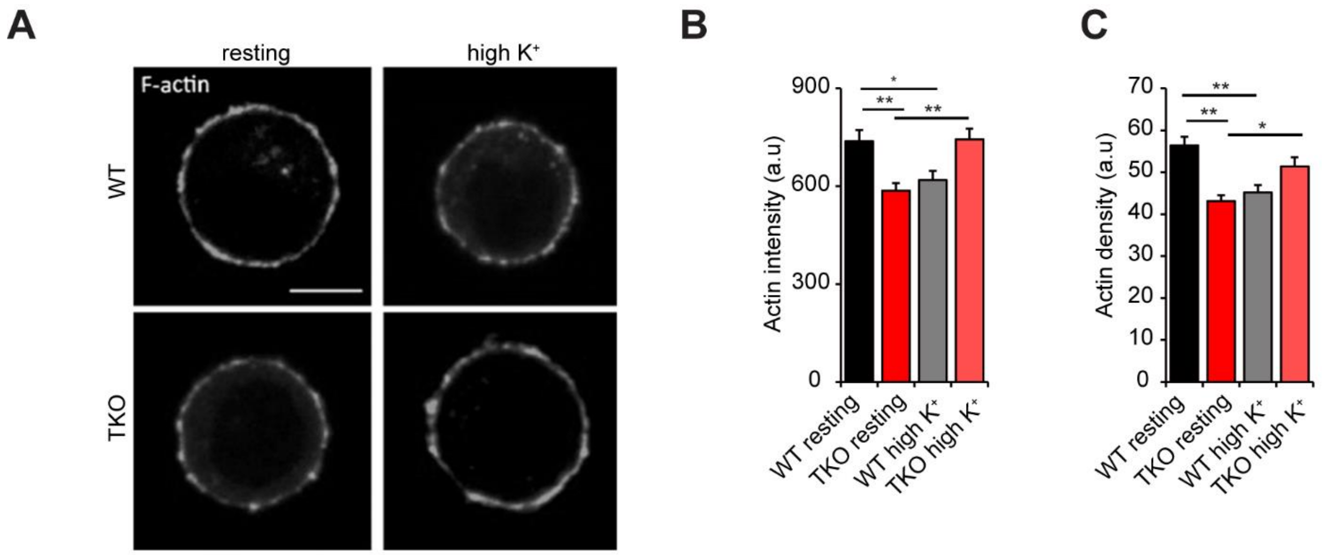

**Suppl. Fig. S8.**

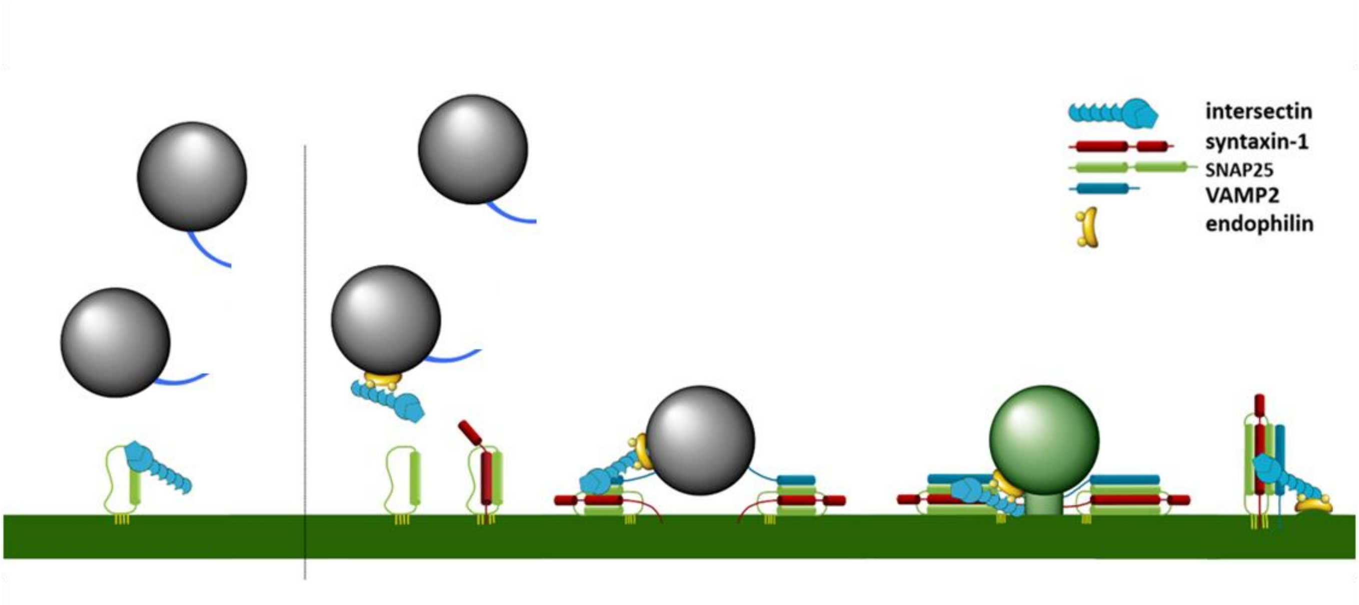
