## Supplemental data_text for "Endophilin-A controls recruitment, priming and fusion of neurosecretory vesicles"

**Supplemental Information**

**Supplemental Material and Methods**

Adrenal chromaffin cells on poly-L-lysine-coated glass coverslips were washed carefully in pre-warmed Locke's solution before incubation in extracellular (control condition) or high $K^+$ solution (88 mM NaCl, 59 mM KCl, 2 mM $CaCl_2$, 1 mM $MgCl_2$, 10 mM HEPES, and 2 mg/ml D-glucose, pH 7.20, 300 mOsM/kg) for 3 minutes at RT. The cells were then fixed in freshly prepared 4% PFA for 30 minutes at RT and neutralized with 50mM $NH_4Cl$ in PBS for 20 minutes. The cells were then permeabilized and blocked for unspecific binding using blocking solution (3% BSA, 0.2% coldfish gelantine, 1% goat serum, 0.1% Triton X-100 in PBS, pH 7.4) at RT. Subsequently, the cells were incubated for 30 min with phalloidin-Texas Red dye for staining F-actin (1U/cs; 200U/mL stock dissolved in methanol), washed, stained with DAPI and mounted using Mowiol-488® mounting medium. The images were acquired using Zeiss LSM 800 laser scanning confocal microscope (63x objective, numerical aperture 1.4). The actin staining was analyzed by ImageJ PLASMaCC macro reported in Kurps et al. (2014). Briefly, the signal in the proximity of plasma membrane was measured by a polar transformation function in ImageJ and the mean intensity and average density was calculated. The data are represented as mean±SEM, and statistics was performed by one-way ANOVA.

The band signal intensity in Western blots were quantified by measuring the mean intensity of ROI in ImageJ software, and subtracting the background signal from the adjacent area.

**Supplemental Figure Legends**

**Figure S1** (related to Figure 1)**. Endophilin on isolated LDCVs and re-distribution of endophilin upon chromaffin cell stimulation**

(A) Scheme of the protocol used to isolate LDCVs from the medulla of bovine adrenal glands. (B) Immunoblot analysis of LDCVs isolated from bovine adrenal medulla and SVs from mouse brain revealed the presence of endophilin 1 at LDCVs and SVs. Syt1 was used as a vesicle marker protein. Same volume (10µl: ~50µg homogenates, ~10µg vesicles) of samples were loaded. (C-D) Upon stimulation with the high $K^+$ solution, distributions of endophilin 1 and endophilin 2 were mildly altered - more endophilin 1 and endophilin 2 were present at the plasma membrane after stimulation, as indicated by the intensity line profiles. Scale bar 2 µm.

**Figure S2** (related to Figure 2)**. Double-log plot shows correlation between the lower time constant of the RRP and the lower pool size of the RRP.**

The RRP was smaller (contained fewer vesicles), but RRP vesicles fused faster in the endophilin TKO chromaffin cells (see Figure 2). We considered the possibility that those two phenotypes are linked. Double-log plot of time vs. the pool size of the RRP showed correlation, and suggested that the lower pool size and the faster time constants of fusion may constitute a single phenotype. A similar observation was made in synaptotagmin-7 KO cells (Schonn et al., 2008).

**Figure S3** (related to Figure 3)**. Expression of endophilin 1 and endophilin 2 rescued exocytosis in endophilin TKO cells (2$^{nd}$ stimulus)**

(A) Verification of lentiviral expression system for endophilin 1 and endophilin 2 in HEK293 cells. Note that the two proteins were expressed to similar levels (quantification is not possible due to the use of two different antibodies). (B) EGFP fluorescence intensities measured from cells expressing endophilin 1 and endophilin 2 using bicistronic lentiviral systems were comparable (50 cells from 3 independent experiments). (C-D) Expression of endophilin 1 and endophilin 2 rescued the exocytic defects seen in endophilin TKO cells (stimulus #2). Panel arranged as in Figure 3A, with 3 groups: endophilin TKO (red traces; mean of 24 cells), endophilin TKO + endophilin 1 (green traces; mean of 23 cells) and endophilin TKO + endophilin 2 (blue traces; mean of 25 cells). Endophilin KOWTKO data from Figure 2 (black trace; mean of 30 cells) were superimposed. Note that endophilin 1 as well as endophilin 2 could rescue exocytosis, but endophilin 2 was more efficient. (E-F) Burst and sustained component were rescued upon expression of endophilin 1 and 2, respectively. (G) Measure of recovery, calculated as the ratio of burst secretion of the second over the first stimulus was not significantly altered.

**Figure S4** (related to Figure 5). **Levels of key exocytic proteins were unaltered in mouse adrenal gland homogenates**

(A) Representative Western blots for chromogranin-A (CgA), synaptotagmin-1 (Syt-1), synaptophysin and SNAP25 showed no difference between total protein levels in endophilin TKO cells and controls (littermate KOWTKO and WT). (B) Quantification of Western blot data shown in A (one-way ANOVA, 3-4 independent experiments were performed). (C-D) Western blots for Munc18-1, syntaxin-1 and VAMP2 showed no difference between total protein levels in endophilin TKO cells and controls (littermate KOWTKO and WT). Quantified in (D) (one-way ANOVA, at least three independent experiments were performed).

**Figure S5** (related to Figure 6). **Characterization of uptake of mCling-Atto647 and chrolar toxin**-B-A594 **endocyctic assays in adrenal chromaffin cells.** (A) WT mouse chromaffin cells were incubated with either negative control for Pitstop-2 (A), or Pitstop-2 (B) for 10 min prior the start of the experiment. Images were acquired at 30s, 2.30 min and 4.30 min after mCling-Atto647 dye addition, stimulation (high $K^+$) was applied 30 s later (shown as 0, 2 and 4 min in the panel). Scale bar 2 um. Considerably lower uptake of mCling-Atto647 was observed in the presence of Pitstop-2, as it can also be seen in Movie 2. (B-C) Endocytic uptake in chomaffin cells was also tested using CT-B-A594 (non-toxic recombinant cholera toxin subunit-B conjugated to Alexa Fluor 594). Scale bar 1.5 µm. Endophilin TKO cells were compared to the littermate controls (endophilin KOWTKO and WT; note that an uptake in only one equatorial optical plane was analyzed, not in the whole cell as in the case of transferrin). Quanified in (C) (min. 20 cells from 2 independent experiemnts, one-way ANOVA).

**Figure S6** (related to Figure 8)**. Levels of intersectin-1 protein were not altered in endophilin TKO cells.**

(A-B) Western blot for intersectin 1 (ITSN-1) showed no difference between total ITSN-1 protein levels in endophilin TKO and controls (endophilin KOWTKO and WT); quantified in (B) (one-way ANOVA, three independent experiments). (C) Immunoblot analysis of LDCVs (as in Figure S1B) against ITSN-1 and ITSN-2 detected the presence of small amount of ITSN-1 and ITSN-2 proteins at LDCVs and SVs. (D-E) Quantification of ITSN-1 distribution in cell cytosol (D) and near the membrane (E) at resting and stimulated (depolarization by high $K^+$) conditions showed altered ITSN-1 distribution in the endophilin TKO cells (mean±SEM; one-way ANOVA). (F-G) Quantification of ITSN-2 (same as ITSN-1) on cell cytosol (F) and near the membrane (G) are represented as bar graphs (mean±SEM; one-way ANOVA). The ratio of membrane to cytosol intensity is shown in Figure 8A-D.

**Figure S7** (related to Figure 8)**. F-actin network is altered in the endophilin TKO cells**

(A) Confocal image (through the equatorial plane) of WT and endophilin TKO chromaffin cells stained with phalloidin-Texas Red at resting vs. stimulated (depolarization by high $K^+$ solution) conditions. Note that the intensity and density of F-actin was lower in the proximity of the plasma membrane in endophilin TKO cell at resting condition. While stimulation in WT cells led to the depolymerisation of the actin barrier and the re-polymerisation of the *de novo* actin filaments at the plasma membrane, the opposite was detected in TKO cells. Scale bar 2 µm. (B-C) Quantification of the cortical F-actin intensity (B) and average density (C) using PLasMACC plugin in ImageJ software (30 cells, one-way ANOVA).

**Figure S8. Model of endophilin and intersectin's role in exocytosis.** Endophilin is present on at least some neurosecretory vesicles and has a role in the vesicle recruitment, priming and fusion. It can also regulate intersectin localization, possibly to ensure that intersectin acts only at the optimal time. (Left) Without endophilin, intersectin mislocalizes to the plasma membrane so its reported role in exocytosis is likely altered in endophilin TKO cells. (Right) We propose that endophilin and intersectin act in tandem to stimulate the recruitment of secretory vesicles to the plasma membrane and to their site of release, expectedly through modulation of the actin network. In addition, endophilin and intersectin may stabilize the SNARE complex between the vesicular VAMP2/synaptobrevin-2 and plasma membrane-resident SNAP25 and syntaxin-1. We also propose that, once recruited to the plasma membrane, endophilin and intersectin do not dissociate after exocytosis but take part in the subsequent endocytic steps. As such, endophilin and intersectin act as a scaffold that couples exocytic and endocytic events.
In addition to data presented here, this model is supported by our previous results on endophilin and intersectin (Milosevic et al., 2011; Pechstein et al, 2015) and a number of other reports (e.g., Okamoto et al., 1999; Malacombe et al., 2006; Yu et al., 2009; Momboisse et al., 2010; Bai et al., 2010; Winther et al., 2013; Sakaba et al, 2013; Gerth et al., 2017).

### MOVIES

**Movie 1**. Wild-type mouse chromaffin cell filmed 30s after addition of mCling-Atto647 dye for 12 minutes. The high potassium (stimulates exocytosis and endocytosis in chromaffin cells) was added after 2 minutes. Movie speed 10f/s.

**Movie 2**. Wild-type mouse chromaffin cell incubated with Pitstop-2 for 10 minutes before mCling-Atto647 dye is added. The cells is filmed 30s after addition of mCling-Atto647 dye for 15 minutes. The high potassium (stimulates exocytosis and endocytosis in chromaffin cells) was added after 3 minutes. Note that the cell membrane surfaces increased since endocytosis was efficiently inhibited. Movie speed 10f/s

**Movie 3**. Endophilin KOWTKO mouse chromaffin cell filmed 30s after addition of mCling-Atto647 dye for 8 minutes. Movie speed 10f/s.

**Movie 4**. Endophilin TKO mouse chromaffin cell filmed 30s after addition of mCling-Atto647 dye for 8 minutes. Movie speed 10f/s.
